# A missense *KCNQ1* Mutation Impairs Insulin Secretion in Neonatal Diabetes

**DOI:** 10.1101/2021.08.24.457485

**Authors:** Zhimin Zhou, Maolian Gong, Amit Pande, Ulrike Lisewski, Torsten Röpke, Bettina Purfürst, Lei liang, Shiqi Jia, Sebastian Frühler, Anca Margineanu, Chun Zeng, Han Zhu, Peter Kühnen, Semik Khodaverdi, Winfried Krill, Wei Chen, Maike Sander, Klemens Raile, Zsuzsanna Izsvak

## Abstract

KCNQ1/Kv7 is a voltage-gated K^+^ channel that regulates heart rhythm, glucose and salt homeostasis. Mutations of *KCNQ1* are primarily associated with long-QT syndrome and type 2 diabetes; however, thus far *KCNQ1* mutations have not been associated with monogenetic diabetes. Here, we identified a homozygous *KCNQ1* missense mutation (R397W) in an individual with permanent neonatal diabetes (PND). To identify the mechanisms that link the mutation to the disease, we introduced the mutation into human embryonic stem cells (hESCs), and used them to derived pancreatic β-like cells (hESC-β cell). In early β-like cells, we observed atypical membrane electrical activity, increased levels of cytoplasmic Ca^2+^, and a hypersecretion of insulin. Upon extended culture, their insulin secretion decreased and the number of apoptotic cells increased, resulting in a reduction in the numbers of β-like cells. Late-stage β-like cells exhibited a decrease in the expression of metabolic genes, e.g. HNF4α, PDX1 and GLUT1, providing a possible mechanism for β-cell dysfunction. Our study identifies *KCNQ1* as a novel candidate gene of monogenetic diabetes and shows that *KCNQ1* regulates β-cell function and survival.

## Introduction

The potassium Voltage-Gated Channel Subfamily Q 1 (KCNQ1) channel plays physiological roles in the cardiovascular system, intestine and pancreas (Neyroud, Tesson et al., 1997, Schroeder, Waldegger et al., 2000, Waldegger, Fakler et al., 1999, Kottgen, Hoefer et al., 1999), where it regulates several essential processes, including repolarization of cardiomyocytes, vasodilatation and secretion (Barhanin, Lesage et al., 1996, Chadha, Zunke et al., 2012, Jepps, Chadha et al., 2011, Zeng, Guo et al., 2016). The KCNQ1/KV7 channel induces voltage-dependent current by rapidly activating and slowly deactivating potassium-selective outward current (Chouabe, Neyroud et al., 1997, Sanguinetti, Curran et al., 1996, Sachyani, Dvir et al., 2014). Hereditary mutations of *KCNQ1* are primarily associated with cardiovascular pathologies (e.g. long QT syndrome 1, short QT syndrome 2, familial atrial fibrillation or Jervell and Lange-Nielsen syndrome) (Valente, Sparago et al., 2019, Chen, Xu et al., 2003, Bellocq, van Ginneken et al., 2004). *KCNQ1* is also a strong candidate for conferring susceptibility to type 2 diabetes (T2D) (MacDonald & Wheeler, 2003, Unoki, Takahashi et al., 2008, Yasuda, Miyake et al., 2008, Chiou, Zeng et al., 2021). So far, however, mutations in *KCNQ1* have not been associated with monogenic diabetes.

Roughly 2-6% of all patients with diabetes have a hereditary monogenic form that arises from one or more gene mutation(s) in a single gene (Thomas, Brackenridge et al., 2016, Cho, Shaw et al., 2018). While MODY summarizes the majority of hereditary monogenic diabetes cases (most common mutations in glucokinase (GCK), hepatocyte nuclear factors (HNF) 1*α*, 4*α*, 1β) (Sanyoura, Philipson et al., 2018), NDM is much rarer, accounted only about 0.012% of all live births (Grulich-Henn et al., 2010). In contrast to maturity-onset diabetes (MODY), which develops in adolescence or early adulthood (Hattersley & Patel, 2017), neonatal diabetes mellitus (NDM) is usually diagnosed within the first 180 days of life and is more severe (Grulich-Henn, Wagner et al., 2010). Patients with transient or permanent NDM exhibit severe β-cell dysfunction accompanied by a reduction in islet cell mass and sometimes even pancreas aplasia (Polak & Cave, 2007, Aguilar-Bryan & Bryan, 2008). If insulin is absent during fetal development, the fetus could fail to thrive after birth. Even in milder cases, neonates with NDM are marked to suffer from life-long acute hyperglycemia and life-threatening dehydration (Polak & Cave, 2007, Mohora & Stoicescu, 2016).

So far NDM has been associated with mutations in the *ABCC8, KCNJ11,* and *INS* genes. Here, we report a permanent NDM (PND) patient with a homozygous *KCNQ1* mutation (C1189T). The mutated KV7 channel protein *KCNQ1*, contributing to permanent NDM is especially interesting. The hereditary mutations of *KCNQ1* are primarily associated with cardiovascular and not from metabolic disorders. Although several lines of evidence supports an association between *KCNQ1* and the regulation of insulin secretion (MacDonald & Wheeler, 2003, Unoki et al., 2008, Yasuda et al., 2008, Rorsman & Ashcroft, 2018), what exact role a KCNQ1/Kv7 channel might play in the glucose-stimulated insulin secretion process is still unclear. Notably, *KCNQ1* is part of an epigenetically regulated (e.g. imprinted) genomic locus (Nakano, Murakami et al., 2006), thus, besides a potential loss of function, the mutation might be also associated with impaired expression regulation. As a strong T2D candidate (MacDonald & Wheeler, 2003, Unoki et al., 2008, Yasuda et al., 2008), *KCNQ1* might represent a potential link between various types of diabetes. The other reason is that the regulation of insulin secretion might work differently in humans and other mammalian species (Asahara, Etoh et al., 2015).

Motivated by the challenge, we sought to characterize mechanisms linking the *KCNQ1* mutation to PND. Recently, the technology of generating disease-relevant human pluripotent stem cell (hPSC)-based organoid models became a vital tool in diabetes research and precision therapies (Zeng et al., 2016, Chiou et al., 2021, Zhou, Kim et al., 2018). We used CRISPR/Cas9 to introduce the *KCNQ1* mutation into hESCs and then derived pancreatic endocrine organoids. The organoids proved suitable to mimic the early events in human pancreatic β-cell differentiation, but also study biological processes at later stages of maturation. The mutant cells were characterized using cellular assays and transcriptome analyses. Our data revealed that the identified (C1189T) mutation does not affect the regulation of expression or pancreatic differentiation. It is a missense mutation (R397W), resulting in a loss of channel function. Although, the mutation was previously reported from an intrauterine death case associated with a long QT syndrome 1 (LQT1), our study shows that the mutation causes an atypical membrane polarization of β-like cells, resulting in a variable insulin secretion phenotype. The detailed characterization of KCNQ1 function during β-cell development allowed us to decipher the role of KCNQ1 in regulating glucose metabolism in a human-specific way. This gives us a better understanding its role in pathology of NDM.

## Results

### Exome sequencing identifies a homozygous missense KCNQ1^R397W^ mutation in a permanent neonatal diabetes patient

Here, we investigate a male patient born with intrauterine growth retardation and no detectable C-peptide in his blood at birth or even later. The patient was diagnosed with permanent NDM (PND) (Fig 1A). Whole-exome sequencing of the patient and his family members revealed four candidate gene variants in *KCNQ1, MIA3, NAXD* (alias *CARKD*) and *MYO1F,* respectively. None of them had been previously reported in neonatal diabetes. From this list, MYO1F is not expressed at significant levels in human islets (Chiou et al., 2021). Studies had shown an association between the three other candidates and cardiovascular diseases (e.g. LQT1 (Ghosh, Nunziato et al., 2006), coronary artery disease (Samani, Erdmann et al., 2007)) or neurological disorders (Van Bergen, Guo et al., 2019, Malik, Nadir et al., 2020). Importantly, dysfunctional *KCNQ1* had been reported to impair glucose stimulated insulin secretion (GSIS) in differentiated β-like cells, in both mice and humans (Zeng et al., 2016, Boini, Graf et al., 2009). *KCNQ1* is also a strong candidate for type 2 (T2D) diabetes susceptibility (Unoki et al., 2008, Yasuda et al., 2008), whereas *MIA3* and *CARKD* are weak candidates (*MIA3*, p value< 1e-4; *CARKD*, p value=2.4e-5) (Aylward, Chiou et al., 2018). Sanger sequencing confirmed the 1189 C>T nucleotide change in the exon 9 of the *KCNQ1* gene, resulting in a missense mutation (R397W) (Fig 1A). The PND patient is homozygous for this mutation, while other members of his consanguineous family carry the mutation in a heterozygous form and are healthy (Fig 1A). Taken together, the mutated *KCNQ1*^R397W^ was the best candidate as the cause of patient’s diabetic phenotype, so we chose it for a follow-up study to decipher its potential contribution.

**Figure 1.**
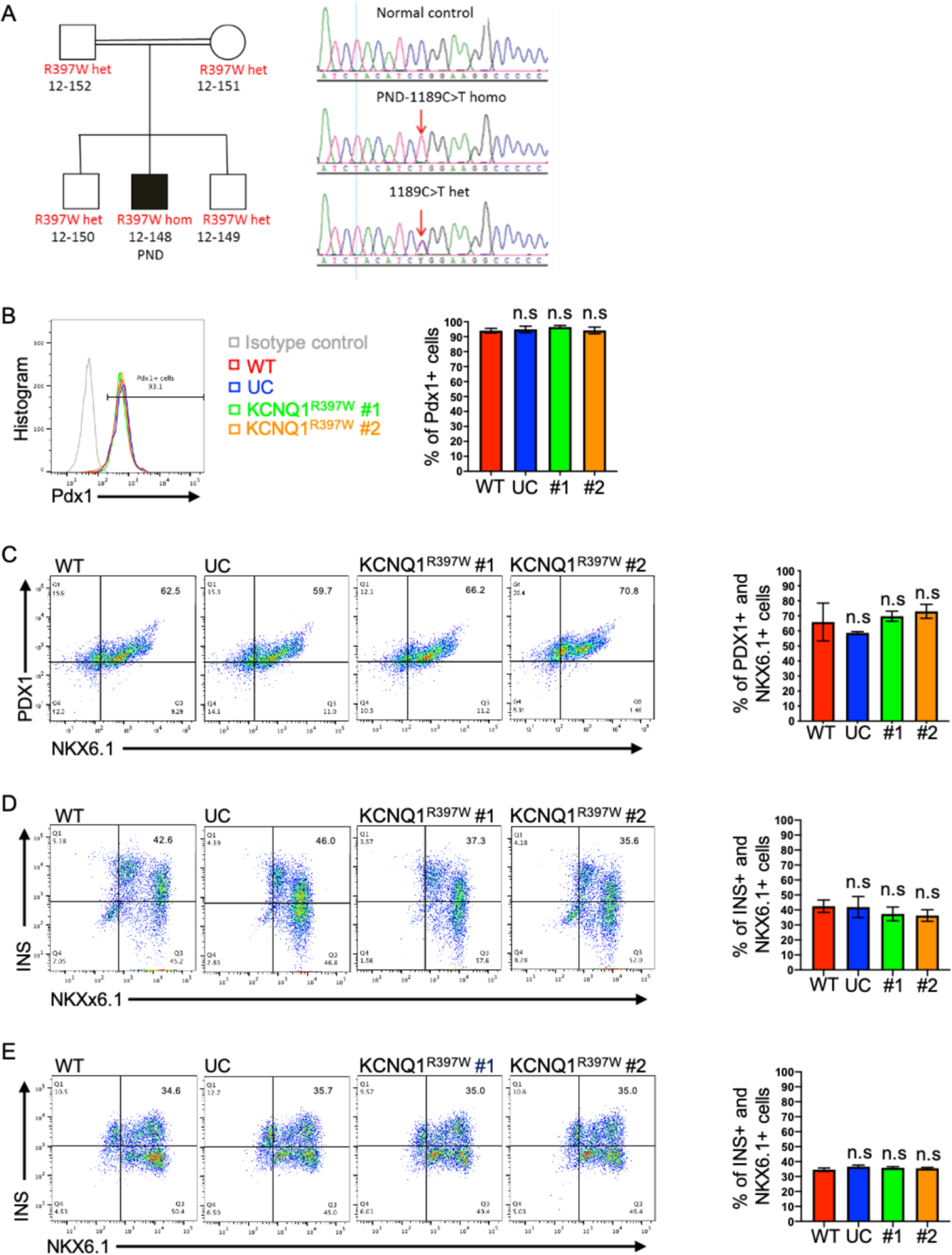
KCNQ1^R397W^ hESCs normally differentiated to human β-like cells. A. Pedigree of the patient’s family. Squares and circles represent males and females, respectively. The patient (marked in black) diagnosed by permanent neonatal diabetes (PND). Other members of his family are healthy. The region of the 1189 C>T mutation in KCNQ1 is shown from control; homozygous (homo) mutation from the patient; patient’s family members (heterozygous). B-E. Flow cytometry analysis and quantification of cells using stage-specific markers. Pancreatic endoderm cells (day 11) express PDX1 (B). Pancreatic endocrine precursor cells (day 14) express PDX1 and NKX6.1 (C). Immature β cells (day 21) express NKX6.1 and insulin (INS) (D). Matured β cells (day 31) express NKX6.1 and insulin (E). Data information: In (B-E), data are presented as mean ± SD. n.s indicates a non-significant difference, *p < 0.05, **p < 0.01, ***p < 0.001, and ****p < 0.0001 (Student’s t-test).

### KCNQ1^R397W^ mutant hESCs differentiate normally into pancreatic β-like cell

To mimic the effects of the 1189 C>T mutation in the patient, we generated an *in vitro* hESC-derived pancreatic islet-like organoids. First, CRISPR/Cas9 engineering was used to generate the homozygous point mutation in hESCs_H1. We established two

*KCNQ1* mutant clones from the CRISPR/Cas9 transfected cell library (KCNQ1^R397W^ No.1 and No.2), whereas wild type (WT) and unmodified control (UC) colonies served as controls (Fig EV1A). Importantly, all of the colonies had identical hESCs morphology, stained positive for SOX2 and OCT4 (Fig EV1B), and expressed levels of OCT4, SOX2 and NANOG pluripotency markers similar to those of wild type (Fig EV1C). This indicated that the editing did not compromise pluripotency.

To determine whether the mutation affected β-cell development, we differentiated the two mutant *KCNQ1* hESCs_H1 clones and their controls into pancreatic insulin+ organoids. Our protocol, based on published methods (Chiou et al., 2021, Velazco-Cruz, Song et al., 2019), was further optimized in-house (see Methods). To monitor the differentiation process, we quantified cells at each stage using flow cytometry and stage-specific markers. PDX1 (Pancreatic duodenal homeobox-1 protein) binds to the promoter of insulin and initiates insulin expression (Wang, Cahill et al., 1999, Wang, Zhou et al., 2001), and is a marker for pancreatic endoderm. As a first step, we determined the percentage of PDX1+ cells (Fig 1B). In the subsequent steps, we quantified PDX1+/NKX6.1+ (NK6 homeobox 6) cells (pancreatic endocrine precursors; Fig 1C); NKX6.1+/insulin+ cells (immature β cells at early stages; Fig 1D); as well as NKX6.1+/insulin+ cells (matured β cells at later stages; Fig 1E). We detected no significant differences between the clones in terms of the composition of cell types or organoid morphology (Fig EV1D), suggesting that the mutation did not significantly affect the differentiation process.

### The KCNQ1^R397W^ mutation overlaps with a CTCF binding site, but does not affect gene regulation during pancreatic differentiation

Epigenetic modifications in the *KCNQ1* imprinted locus have been reported to affect β cell mass in both human and mice (Asahara et al., 2015, Ou, Yu et al., 2019). As the *KCNQ1* C1189T mutation constitutes a potentially methylated cytosine and overlaps with a predicted motif of the CCCTC-binding factor (CTCF) (Fig EV2A), we asked whether the mutation affected epigenetic gene regulation. Our data mining revealed that this particular CTCF binding site is involved in establishing chromatin loops and regulating gene expression in several human cell lines (e.g. GM12892, human lymphoblastoid and K562, human myelogenous leukemia) (Fullwood, Liu et al., 2009, Sabo, Hawrylycz et al., 2004, Consortium, 2012). If the mutated nucleotide is involved in epigenetic gene regulation, we would expect altered gene expression levels of *KNCQ1* and/or the neighboring genes, resulting in reduced β cell numbers.

To determine whether this particular CTCF binding site is utilized during pancreatic differentiation, we analyzed active enhancer signals (H3K27ac) in ChIP-seq data, which were generated from human cells at different stages of β-cell differentiation *in vitro* (Xie, Everett et al., 2013). We also mined Hi-C data from human islets (Miguel-Escalada, Bonas-Guarch et al., 2019). Our analyses identified dynamic enhancer signals in intron 11 of *KCNQ1* but not in exon 9, in which the mutated locus resides (Fig EV2B). The analysis of the Hi-C data further indicated that the mutated CTCF motif was not part of the pancreatic differentiation program in humans (Fig EV2C). To analyze the genomic methylation status of the mutation site of *KCNQ1*, we performed sodium bisulfite conversion and sequencing using wild type and KCNQ1 mutant cells collected at different stages of differentiation. The assay confirmed that the C1189T mutation abolished methylation, and revealed that this cytosine was a stably methylated cytosine during pancreatic differentiation (Fig EV2D).

Analysis of RNA-seq data of pancreatic differentiation (Xie et al., 2013) revealed that the expression of *KCNQ1* gradually increases after the pancreatic endoderm stage to peak in islets (Fig EV2E). To examine whether the lack of methylation due to the C1189T mutation affected KCNQ1 expression, we performed both RT-qPCR and Western blotting for KCNQ1 at day 31 of differentiation. However, we observed no mutation-associated expression changes (Fig 2A and B). Furthermore, the overlapping regulatory long non-coding (lnc)RNA, *KCNQ1OT1* (Asahara et al., 2015) and the other members of the imprinted locus either exhibited similar levels of expression in all the samples (e.g. *CDKN1C* and *SLC22A18*) or expressed below the threshold of detection (e.g. *PHLDA2*) (Fig 2C).

**Figure 2.**
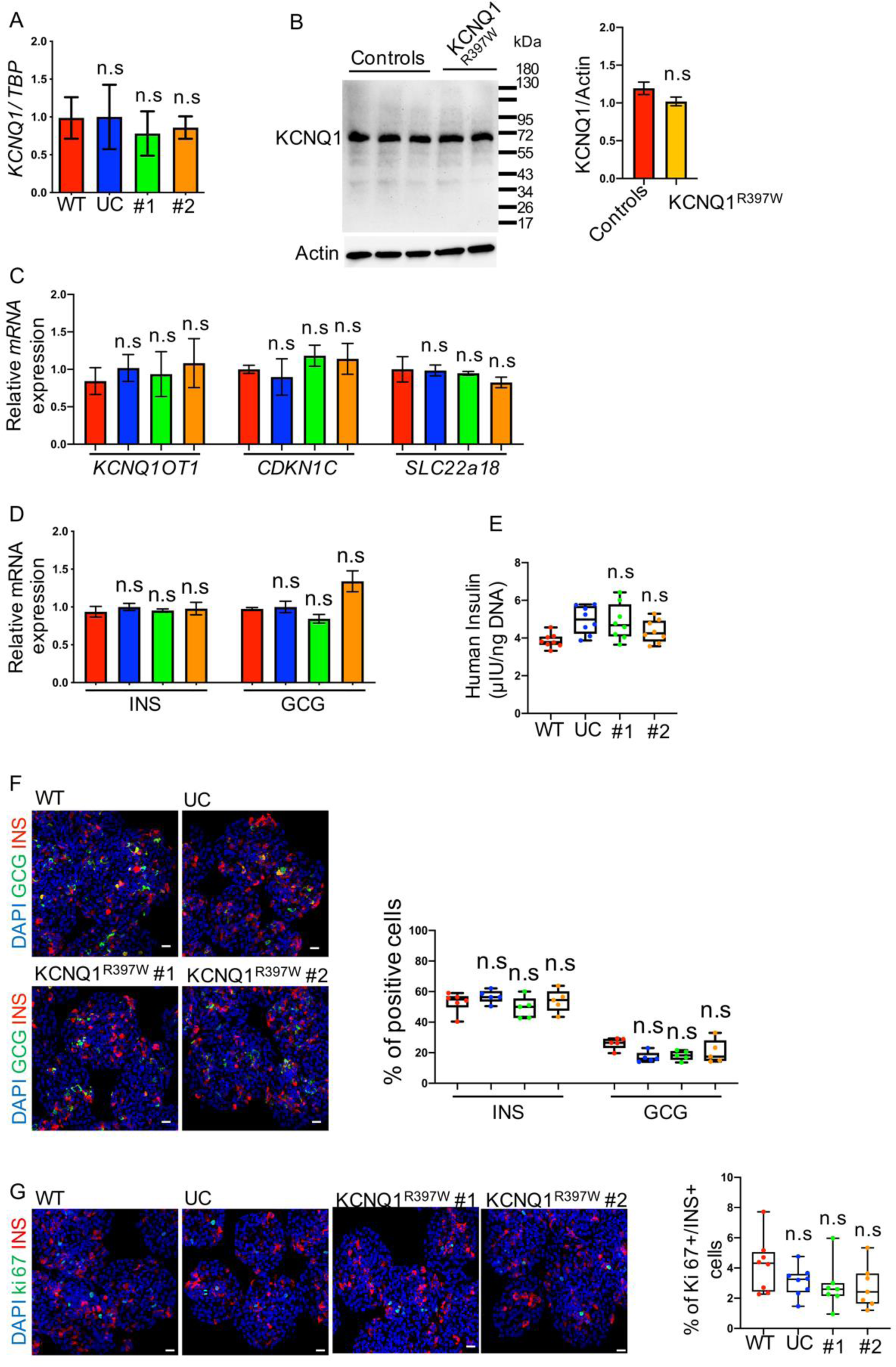
The 1189 C>T mutation did not affect regulatory genomics. A. The *KCNQ1* mRNA level was analyzed by qRT-PCR in human day 31 organoids of WT, UC, and KCNQ1^R397W^. B. Western blot analysis and quantification of KCNQ1 expression in human day 31 organoids of controls (WT and UC) and KCNQ1^R397W^. Data are normalized to ACTIN. C. q-PCR analysis of the expressions of *KCNQ1OT1*, *CDKN1C*, and *SLC22a18* at human day 31 organoids. D. Pancreatic islet-like organoids from WT, UC, and KCNQ1^R397W^ were characterized by qRT-PCR, using glucagon^+^ (GCG+) and insulin^+^ (INS+) as α- and β cell-specific markers, respectively. E. Total insulin content per 1ng DNA of insulin+ cells between KCNQ1^R397W^ and controls matured β-like cells (day 31). F. Immunostaining and quantification INS+ and GCG+ cells at day 31 (matured β cells). G. Immunostaining and quantification of Ki67+ in β-like cells. Scale bar=20 μm. qPCR data are normalized to housekeeping gene *TBP* (TATA-Box Binding Protein). Data information: Scale bar=20 μm (F and G). Data are presented as mean ± SD. n.s indicates a non-significant difference, *p < 0.05, **p < 0.01, ***p < 0.001, and ****p < 0.0001 (Student’s t-test).

Next, we examined whether the C1189T mutation affected the differentiation of pancreatic endocrine cells, including β cells. We analyzed expression levels of pancreatic hormones and quantified hormone+ cells in the pancreatic islet-like organoids. We found no significant differences in the expression of α- and β cell-specific markers (glucagon and insulin, respectively) between mutated and control cells (Fig 2D and E). Percentages of glucagon+ and insulin+ cells were also comparable in the case and control pancreatic islet-like organoids (Fig 2F). Consistent with these findings, the proliferation rates of β-like cells were also similar in mutant and control cells (2-7% Ki67+ cells, Fig 2G). This indicated that in line with the lack of change in the expression regulation of *KCNQ1*, no mutation-associated changes were detected in β-cell numbers or insulin level. Collectively, methylation, ChIP-seq, Hi-C and gene expression analyses suggested that the PND phenotype, caused by the 1189 C>T mutation could not be attributed to changes in gene regulation, thus another explanation was needed.

### KCNQ1^R397W^ β-like cells have a variable insulin secretion phenotype, depending on their maturation state

Alternative to altering gene regulation, the mutation might compromise protein function(s). Besides production, the key function of β cells includes insulin secretion, thus we asked whether the mutation modified this process. To this end, we subjected the organoids to both glucose (GSIS) and KCl depolarization (KSIS) challenges, and monitored insulin secretion at different maturation stages of the β-like cells: *maturing* (day 28), *matured* (day 31) stages. In addition, we included a *late matured stage* after the differentiated cells had been cultured for an extended period (day 40).

While no difference was seen in *KCNQ1* transcript levels between case and control at any of the stages (Fig EV3A), We did, however, noted an altered response of KCNQ1^R397W^ β-like cells to either GSIS or KSIS at both *maturing* (Fig EV3B) and *matured* stages (Fig 3A-D). For example, similar to the low glucose stimulation (LG) in *matured* β-like cells, the controls responded with 7.53±0.515- and 11.04 ±0.995-fold induction of insulin secretion in the KSIS assay. In contrast, the response to the stimulation was significantly higher in the KCNQ1^R397W^ β-like cell lines, yielding 16.87 ± 1.193 and 19.57 ± 0.754-fold enhancement (Fig 3C and D). These observations suggested that the mutant cells secreted elevated levels of insulin at both *maturing* and *matured* stages. Curiously, by contrast, the *late matured* KCNQ1^R397W^ β-like cells had a lower insulin secretion rate than controls in response to glucose stimulation, but not to KCl depolarization (Fig 3E-H). This is interesting, because the insulin content was not significantly different in these cells (Fig 3I), indicating that the change in secretion could not be explained by the altered amount of insulin. Collectively, the mutant β-like cells exhibited a variable, stage-dependent insulin secretion phenotype, with an unusually high levels of secretion at *maturing/matured*- stages, and reduced levels following long-term culturing.

**Figure 3.**
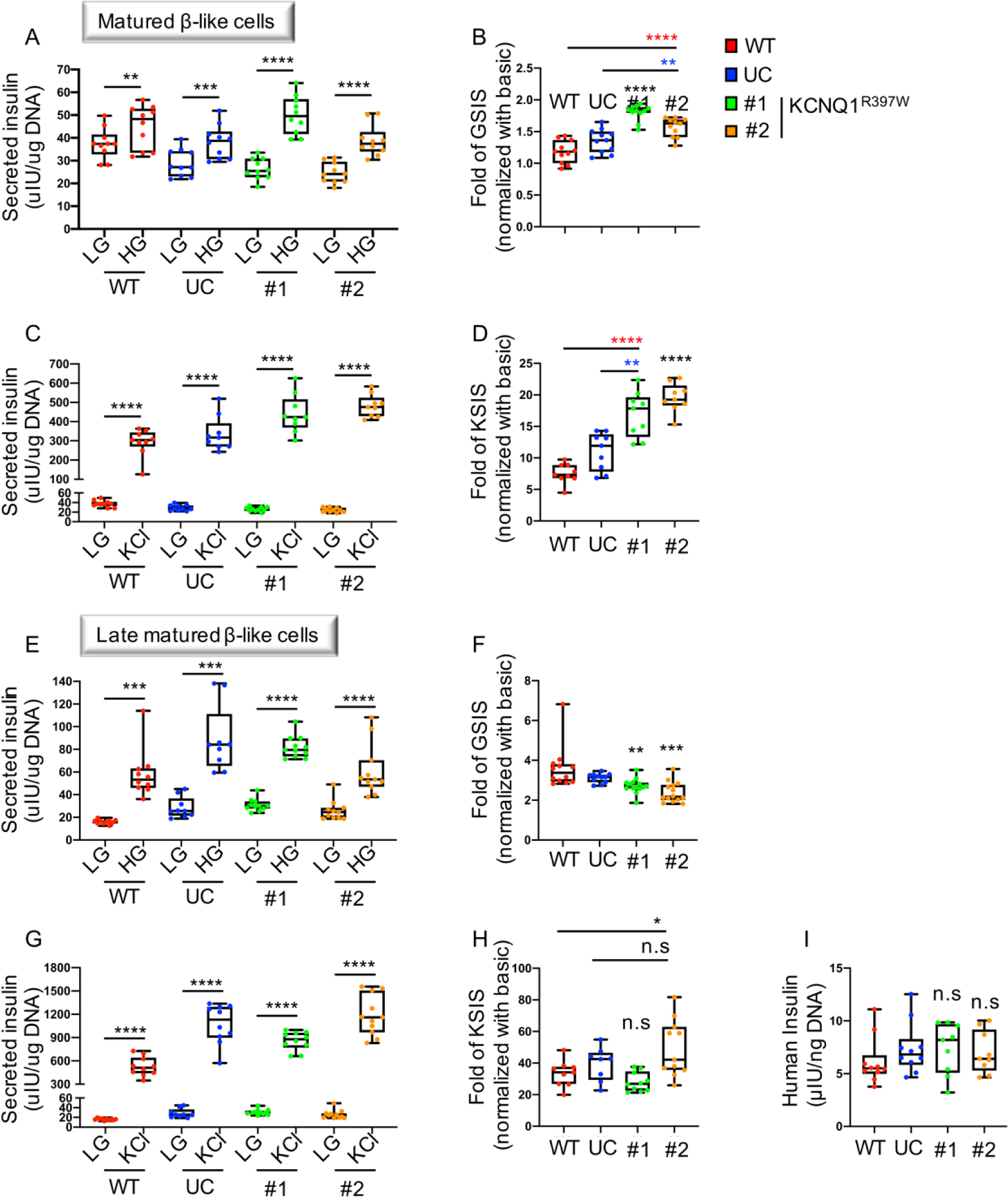
Time/stage-dependent insulin secretion of human β-like cells (day 31 or day 40) upon various challenges, derived from KCNQ1^R397W^ #1, #2, unmodified hESCs (UC) and WT hESCs (WT). A-H. the content of insulin secretion per 1ng DNA of insulin+ cells (Left panels). fold-change of insulin secretion upon various stimulations (Right panels). Day 31 (A and B) and day 40 (E and F) human β-like cells upon 16.8 mM glucose stimulation (GSIS). Day 31 (C and D) and day40(G and H) human β-like cells upon 30 mM KCl stimulation (KSIS). I. Total insulin content per 1ng DNA of insulin+ cells between KCNQ1^R397W^ and controls late-stage β-like cells (day 40). Data information: Data are presented as mean ± SD. n.s indicates a non-significant difference, *p < 0.05, **p < 0.01, ***p < 0.001, and ****p < 0.0001 (Student’s t-test).

To monitor insulin secretion in another model, we also performed the GSIS and KSIS challenges on a mouse immortalized pancreatic β-cell line (referred to as SJ β cells (Jia, Ivanov et al., 2015)), which had been transfected with KCNQ1^R397W^ or WT expression constructs. Similar to our findings in the human model, SJ β cells, which expressed the mutant *KCNQ1* had an altered insulin secretion rate in response to both challenges (Fig EV3C). However, we noticed two differences. First, we did not observe an increase in insulin secretion in the mouse model; it decreased upon both challenges. Second, the total insulin level was already lower in the KCNQ1^R397W^ transfected cells compared to WT (Fig EV3D). This suggested that the reduction in insulin secretion in mouse β cells might be explained by changes in overall insulin expression, which was not the case for the human cells. This suggests that the reduced insulin secretion, by contrast to mutant human β-like cells, might also be explained by altered insulin expression in mouse β cells. Thus, while the mutation ultimately affected insulin secretion in both models, the mechanisms causing this might differ between species.

### The KCNQ1^R397W^ accelerates the rhythm of membrane depolarization/ repolarization in human β-like cells

Our next step in investigating potential impairments in the function of KCNQ1^R397W^ was to analyze the structure of the mutant protein. An alignment of the KCNQ1 sequence across species shows that the region immediately surrounding the mutation is highly conserved (Fig EV4A), suggesting that the mutated amino acid might be essential for proper function. In line with this, the PolyPhen2 algorithm predicted that the mutation is “probably damaging” with the high score of 0.918 (Fig EV4B). Our secondary structure analysis predicts that the missense mutation (R397W) of the KCNQ1 alters the structure of Helix A (Fig EV4C), one of the four predicted helical regions (Wiener, Haitin et al., 2008). As this Helix A is implicated in KCNQ1’s protein-protein interactions (Sun & MacKinnon, 2017), the missense mutation of R397W might result in a loss of function of the KV channel.

To test this hypothesis, we investigated membrane polarization. First, we performed Patch-clamp studies in cultured CHO cells, transfected with expression constructs, encoding KCNQ1^R397W^ or KCNQ1^WT^, and recorded KCNQ1 current traces. Compared to WT cells, the current density in KCNQ1^R397W^ expressing cells was reduced (Fig 4A), consistent with effects on membrane polarization. Curiously, another patch-clamp study of HEK293 cells with the same R397W mutation, identified from a LQT1-associated fetal death case *in utero*, had produced similar results (Crotti, Tester et al., 2013). This raised questions about how the channel function might specifically be disrupted in β cells, and in the context of glucose metabolism. To pursue this, we also recorded the electrical activity of human *matured* β-like cells that were subjected to GSIS. In addition to the high glucose challenge (20mM), we also used Chromanol 293B (10 μM) treatment (Fig 4B), an inhibitor that blocks KCNQ1/Kv7 channel (Busch, Suessbrich et al., 1996, Bleich, Briel et al., 1997). As expected, in the UC control, both the high glucose concentration and the presence of Chromanol 293B inhibitor elevated the frequency of action potential firing (Fig 4B). In contrast, the frequency of action potential firing was higher in unstimulated KCNQ1^R397W^ pancreatic β-like cells (e.g. at low glucose concentration and in the absence of the inhibitor) (Fig 4B). This observation suggested that the KCNQ1^R397W^ β-like cells have a dysfunctional KCNQ1/Kv7 channel, which leads to an acceleration in the rhythm of membrane depolarization/repolarization in β-like cells.

**Figure 4.**
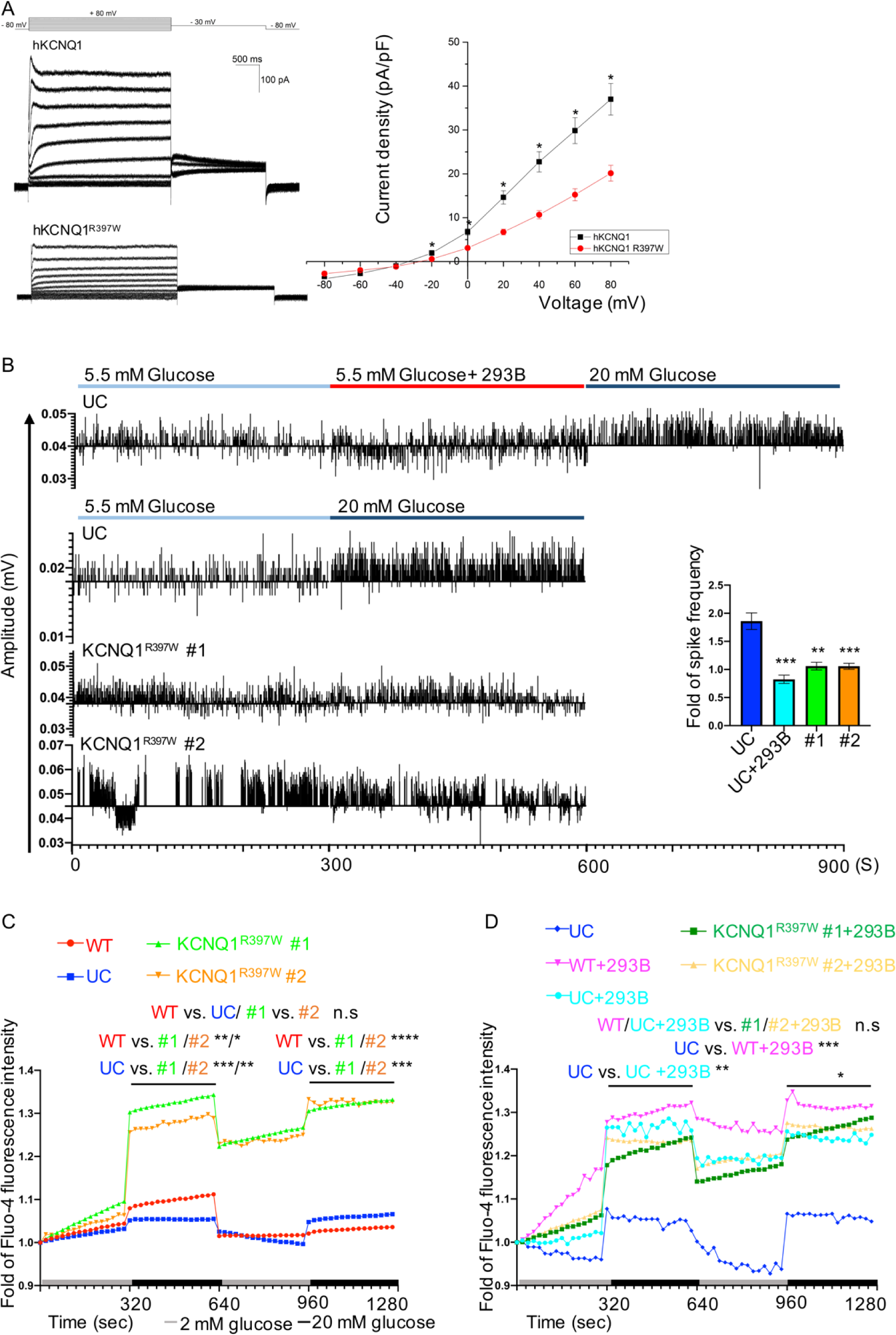
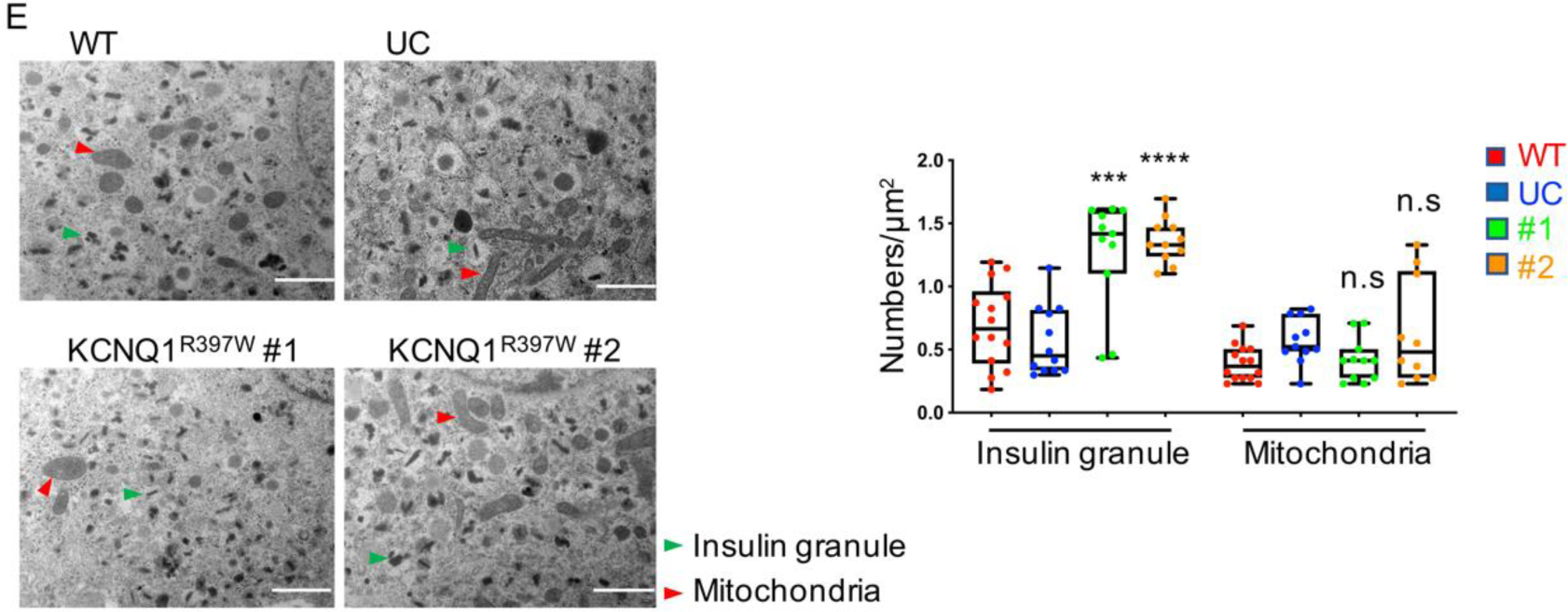
KCNQ1^R397W^ enhances channel activities in β-like cells that results in a cytoplasmic Ca^2+^ accumulation. A. Current traces for KCNQ1^WT^ and KCNQ1^R397W^ in transfected KCNQ1-null Chinese Hamster Ovary (CHO) cells (patch clamp). Data presented as mean ± SEM, p values calculated by two-way ANOVA. B. Recording of electrical activity and quantification of the spike frequency of human mature β-like cells induced by 5.5 mM glucose with/without 10 μM Chromanol 293B (293B) and 20 mM glucose. Data presented as mean ± SD (Student’s t-test). C, D. The dynamic Ca^2+^ flux analysis of β-like cells by Fluo-4 AM staining. β-like cells were cultured with (D) or without Chromanol 293B (293B) (C). Data presented as mean; p values calculated by two-way ANOVA. E. Electron microscopy images and quantification of the crystallized insulin granules (green arrows) and mitochondria (red arrows). Scale bar=1 μm. Data presented as mean ± SD (Student’s t-test). Data information: n.s indicates a non-significant difference, *p < 0.05, **p < 0.01, ***p < 0.001, and ****p < 0.0001.

The proteins of KCNQ1/Kv7 channel needs to traffic to the plasma membrane. As C-terminal *KCNQ1* mutations had previously been reported to alter the trafficking of KCNQ1/Kv7 to the plasma membrane (Wilson, Quinn et al., 2005, Schmitt, Calloe et al., 2007), we monitored the localization of the WT and mutant proteins. Immunofluorescence microscopy revealed that both KCNQ1^WT^ and KCNQ1^R397W^ proteins were similarly localized at the membrane of transfected CHO cell (Fig EV4D), arguing against disrupted trafficking of the mutant form. In addition, 3D confocal scanning microscopy of the human pancreatic β-like cells similarly detected KCNQ1^WT^ and KCNQ1^R397W^ proteins on the membrane in close association with cytosolic insulin (Fig EV4E).

### KCNQ1^R397W^ results in the accumulation of cytosolic Ca^2+^ in human β-like cells

Calcium signaling is involved in synchronizing the periodic change of glucose concentration in β cells (Pagliuca, Millman et al., 2014). To find out whether the elevated channel activities affected Ca^2+^ levels in the mutated β-like cells, we monitored the dynamics of cytoplasmic Ca^2+^ concentrations. In the Fluo-4AM-labelled Ca^2+^ flux analysis, we challenged the mutant and control organoids in their *matured* stage with either high glucose stimulation (GSIS) or chromanol 293B inhibitor treatment. The analyses revealed increases in the cytoplasmic Ca^2+^ concentrations in both control and mutant β-like cells in response to the high glucose stimulation (Fig 4C). However, the mutant cells had a relatively more robust response to the elevated glucose stimulation, and did not return to the baseline in their “resting” stage (Fig 4C). Similar, under chromanol 293B treatment, the mutant cells displayed higher elevation of cytoplasmic Ca^2+^ concentrations, and did not completely recover their original levels following the challenge, compared to cells that were not treated with inhibitors e.g. WT, UC and human islets) (Fig 4D and EV4F). The observation that the mutant cells had a more robust response to stimulation and failed to properly return to their baseline suggested that these cells might eventually accumulate an abnormally elevated cytosolic Ca^2+^ level upon challenges with glucose.

### KCNQ1^R397W^ β-like cells accumulate crystallized insulin granules

How would elevated levels of cytosolic Ca^2+^ affect β cells? In β cells, Ca^2+^ contributes to the formation of secretory granules that structurally organize insulin in (Zn^2+^)2(Ca^2+^)Insulin6 crystals (Rorsman & Ashcroft, 2018, Dunn, 2005). We used electron microscopy to determine whether the insulin granules differed between mutant β-like cells and their controls. We observed that the structure of insulin granules was consistent with their *matured* stage, and was similar to those reported from human islets (Deconinck, Potvliege et al., 1971). However, β-like cells derived from the mutant clones had a significantly higher number of crystallized insulin granules than the controls (e.g. wild type and unmodified control) (Fig 4E). These data suggested that the elevated cytoplasmic Ca^2+^ level resulted in a more extensive crystallization of insulin, leading to an accumulation of secretory granules. As a downstream effect, this might be linked to the observed elevation in the rate of insulin secretion.

### Reduced glucose transporter 1 (GLUT1) expression in *late matured* KCNQ1^R397W^ β-like cells

Could the dysfunctional KCNQ1/Kv7 channel disturb global gene expression and affect biological processes in human β-like cells? To answer, we performed RNA-sequencing on mutant and control samples (at day 32, *matured* stage) and compared their transcriptomes. The analysis considered only differently expressed genes (DEGs) that were found in both mutated colonies and were also differently expressed from both controls (e.g. WT and UC) (Fig EV5A). The gene ontology (GO) categories that were most significantly affected included *oxidative phosphorylation*, *MAPK signaling* pathways and *MODY* (Fig 5A).

**Figure 5.**
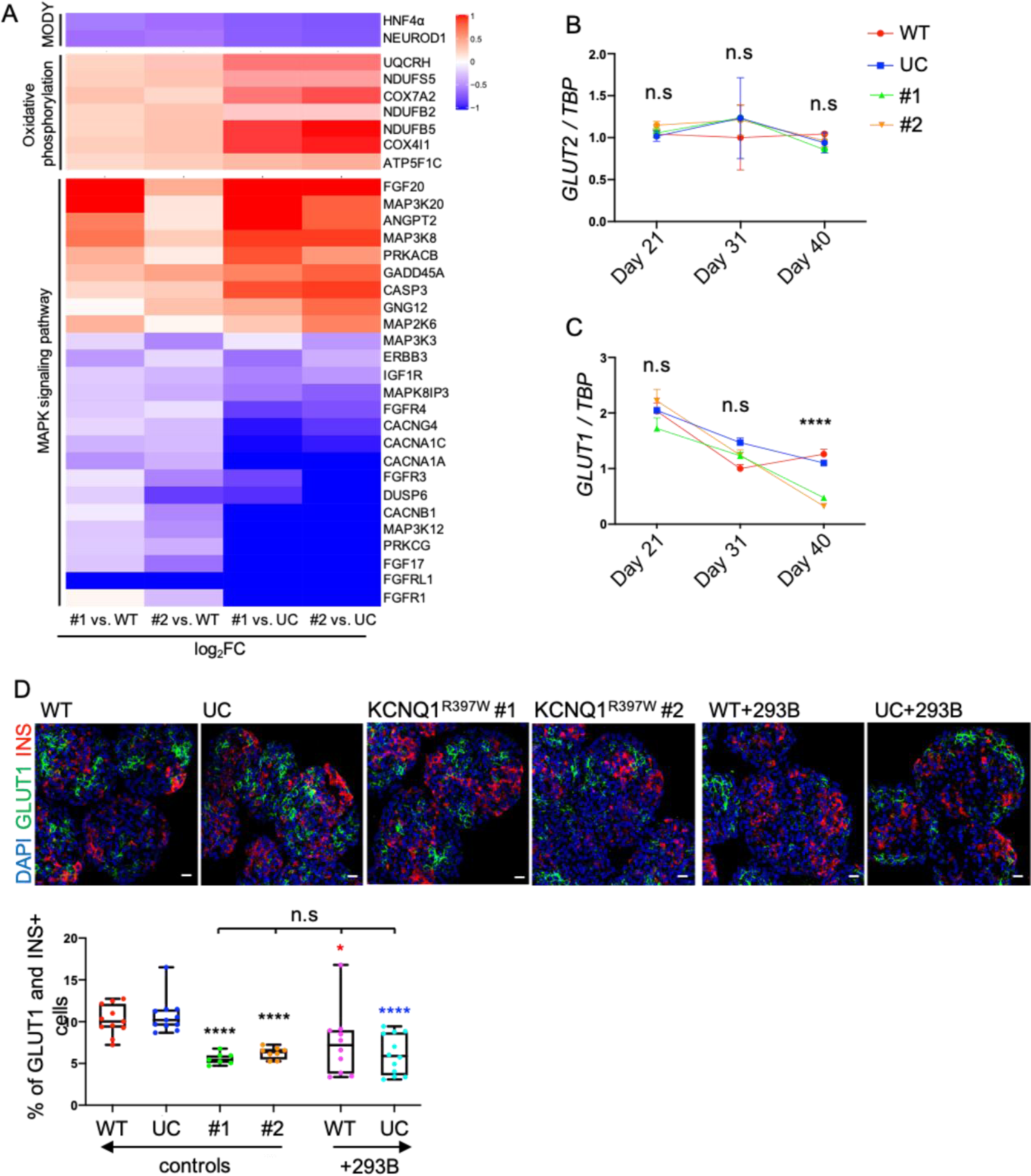
The KCNQ1^R397W^ mutation results in a reduced glucose transport of β-like cells at their late stage. A. Heat map showing the differentially expressed genes (DEGs) revealed by RNAseq analysis. The DEGs belong to the GO categories of *MODY*, *oxidative phosphorylation* and *MAPK signaling*. B, C. Expression qRT-PCR analyses of *GLUT2* (B) and *GLUT1* (C) in stage-specific pancreatic organoids of WT, UC, and KCNQ1^R397W^. Data are normalized to *TBP*. D. Immunoassaying of GLUT1 (green) and insulin (INS, red) positive cells in *late mature* β-like cells (day 40), cultured in normal S7 media (control) or S7 media supplemented with chromanol 293B (+293B) for 12 days (from day 28 to day 40). The INS+/GLUT1+ cells were quantified. Scale bar=20 μm. Data information: The data are presented as mean ± SD. Two-way ANOVA (B and C) or Student’s t-test (D). n.s indicates a non-significant difference, *p < 0.05, **p < 0.01, ***p < 0.001, and ****p < 0.0001.

RNA-seq identified a set of upregulated genes (e.g. *UOCHR, NDUFS5, COX7a, COX411, NDUFB2, NDUFB5, ATP5F1C*) in the *oxidative phosphorylation* pathway (Fig 5A). The upregulation of this metabolic process is expected to increase ATP synthesis in the mitochondria (Siengdee, Trakooljul et al., 2015, Hwang, Lynn et al., 2015, Ikeda, Shiba et al., 2013), which is required for the KATP channel-dependent secretion of insulin by β cells (Haythorne, Rohm et al., 2019, Fu, Gilbert et al., 2013). Electron microscopy revealed comparable numbers of mitochondria in both controls and mutant colonies (Fig 4E), suggesting that the altered oxidative phosphorylation properties of the mutant cells did not affect the mass of these organelles. In the mutant cells, we also observed the transcriptional downregulation of genes upstream of the classical *MAPK signaling* pathway (Fig 5A), including those encoding calcium channel subunits (CACNs) (e.g. *CACNA1A*, *CACNA1C*) and receptor tyrosine kinases (RTKs) RTKs include the members of the fibroblast growth factor receptor (FGFR) family (e.g. *FGFRL1*, *FGFR1*, *FGFR3* and *FGFR4)* that play roles in regulating metabolic features of β cells (Rorsman & Ashcroft, 2018, Wente, Efanov et al., 2006, Miralles, Czernichow et al., 1999, Hart, Baeza et al., 2000). In addition, our RNA-seq analysis identified two down-regulated DEGs: *HNF4α* (*hepatocyte nuclear factor 4α*) and *NEUROD1* (*Neuronal Differentiation 1*) of the MODY gene set of hereditary monogenic diabetes (Fig 5A). *HNF4α* is a transcription factor expressed in liver, kidney and pancreas (Sladek, Zhong et al., 1990, Miquerol, Lopez et al., 1994), and is thought to be a downstream target of FGFRs (Twaroski, Mallanna et al., 2015).

The identification of metabolic genes among the DEGs in the transcriptome of KCNQ1^R397W^ β-like cells suggested a potential link between the metabolic features and the changes in the insulin secretion phenotype (Fig 3). To investigate this further, we generated an expanded list of candidate genes including FGFRs and some of their downstream targets, and performed qPCR on samples collected at the different stages. In the *late matured* KCNQ1^R397W^ β-like cells, qPCR revealed lower levels of *FGFRL1*, *FGFR1* and *HNF4α* than both controls (Fig EV5B). These results suggested a negative effect on metabolic regulation of mutant β-like cells that had been subjected to extended culturing. In a similar assay, we also tested PDX1, which is regulated by *FGFR1* and *HNF4α* (Hart et al., 2000, Narla, Lee et al., 2017, Kuo, Conley et al., 1992, Gerrish, Cissell et al., 2001) (Fig EV5B). We also assayed GLUT2, the major glucose transporter in mice, whose expression level is controlled by *Fgfr1* or *Pdx1* in mouse β-cells (Hart et al., 2000, Waeber, Thompson et al., 1996, Ahlgren, Jonsson et al., 1998, Brissova, Shiota et al., 2002), and which has been previously linked to transient neonatal diabetes (Sansbury, Flanagan et al., 2012). Finally, we added GLUT1 to the list, which has been proposed as the predominant glucose transporter in human β cells (De Vos, Heimberg et al., 1995). Our qPCR analysis revealed a significant expressional decrease of PDX1 and GLUT1 in *late matured* KCNQ1^R397W^ β-like cells, a finding that we also confirmed by immunohistochemical staining (Fig EV5B, EV5C and 5C). However, unlike in mouse β cells, the downregulated *FGFR1* or *PDX1* did not affect the transcription level of *GLUT2* (Fig 5B). This observation can be explained by the fact that each species relies mainly on a different glucose transporters (De Vos et al., 1995). To follow up on the data of *GLUT1*-downregulation in *late matured* stage, we performed additional immunohistochemical staining of GLUT1+ cells, both with and without chromanol 293B inhibitor treatment (Fig 5D). The treatment decreased the number of control GLUT1+ cells to a similar level of the comparator of mutant cells, which had not been treated with the inhibitor (Fig 5D), arguing for a correlation between KCNQ1/Kv7 dysfunction and the expression of GLUT1.

### Chronic exposure to high glucose results in the loss of KCNQ1^R397W^ β-like cells

Because hyperglycemia accelerates β cell dysfunction in type 2 diabetes (Robertson, Harmon et al., 2003), we asked whether the KCNQ1^R397W^ β-like cells exhibited the similar phenotype of glucose-induced toxicity. To this end, we incubated *matured* KCNQ1^R397W^ and control β-like cells for seven days in regular S7 media (control) and in parallel, in S7 media supplemented with high concentration of glucose (20 mM). In high glucose conditions, both flow cytometry and immunostaining revealed a significant decrease in the number of insulin+ KCNQ1^R397W^ β-like cells compared to controls (Fig 6A and EV5D). In regular S7 media, there was a much smaller decreasing trend (Fig 6A and EV5D). This suggests that under chronic exposure to high glucose, the number of insulin+ mutant cells fell at an accelerated rate.

**Figure 6.**
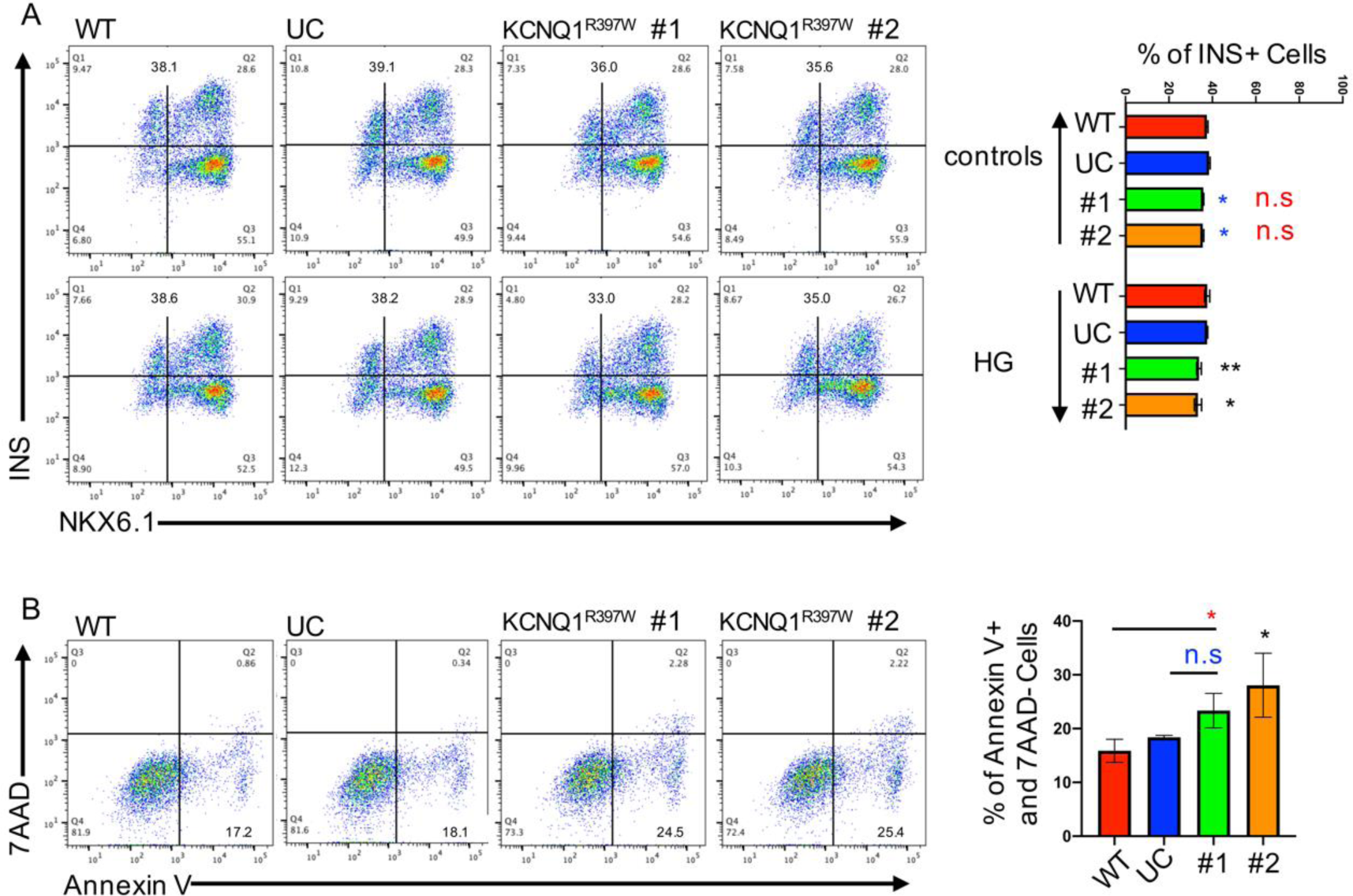
Late-stage KCNQ1^R397W^ β-like cultures harbor pro-apoptotic cells. A. Flow cytometry analysis and quantification of insulin expressing β-like cells. Matured β-like cells were cultured in normal S7 media (controls) or in S7 media supplemented with 20 mM glucose (high glucose, HG) for 9 days (from day 32 to day 40), and subjected to flow cytometry. B. Flow cytometry analysis and quantification of the pro-apoptotic (Annexin V+/7AAD-) cells in late mature β-like cultures. Data information: The data are presented as mean ± SD; n.s indicates a non-significant difference, *p < 0.05, **p < 0.01, ***p < 0.001, and ****p < 0.0001 (Student’s t-test).

Long-term elevation of cytoplasmic Ca^2+^ or overstimulation of the Ca^2+^ signaling pathway have been shown to decrease β-cell mass and function via induced cell death (Bernal-Mizrachi, Cras-Meneur et al., 2010, Kato, Oya et al., 2008, Epstein, Overbeek et al., 1989). To find out whether the accumulated cytoplasmic Ca^2+^ might induce apoptosis at the *late matured* stage of KCNQ1-mutant β-like cells, we quantified cells labeled with pro-apoptotic markers (Annexin V+/7-AAD-) (Fig 6B). FACS quantification revealed a significantly higher number of cells stained for the pro-apoptotic markers in KCNQ1^R397W^ β-like cells at their *late matured* stage, when compared to controls (Fig 6B). This observation pinpoints that induced cell death may contribute to the reduced insulin secretion of KCNQ1^R397W^ β-like cells at their *late matured* stage.

## Discussion

Here, we report the case of a patient, diagnosed with permanent neonatal diabetes (PND). The patient exhibited no detectable endogenous insulin secretion after birth, and carried a homozygous missense C1189T mutation (R397W) of *KCNQ1.* We replicated the defect in a pancreatic organoid model, using CRISPR/Cas9 editing of human embryonic stem cells (hESCs). Using our model, we show that the mutation results in a compromised function of KCNQ1/Kv7 channel, and contributes to a differentiation stage-dependent, variable insulin secretion phenotype. This model also helped us to decipher how the blocked KCNQ1/Kv7 channel could eventually lead to a hypo-insulinemic phenotype of the patient.

In addition to the *KCNQ1* gene, the KCNQ1 locus encodes the lncRNA (KCNQ1OT1), which controls multiple genes in the imprinted genomic locus (Zhang, Zeitz et al., 2014). This epigenetically sensitive genomic region provides a binding platform for several transcription factors including CTCF, which has been implicated in chromatin looping (Ong & Corces, 2014). The C1189T mutation abolishes the methylation of a cytosine and overlaps with a CTCF binding site that is occupied in several tissues (Fullwood et al., 2009, Sabo et al., 2004, Consortium, 2012). 3D chromatin analyses predict that this genomic region can adopt several alternative chromatin-loops that potentially affect gene regulation and are likely regulated on development/tissue specific manner (Sams, Nardone et al., 2016, Tang, Chen et al., 2011, Soshnikova, Montavon et al., 2010, Welsh, Kwak et al., 2015). However, the CTCF affected by the mutation is not used in pancreatic differentiation, and the mutation does not affect the regulation of *KCNQ1* expression on an epigenetic level. Correspondingly, the C1189T mutation does not alter the differentiation and proliferation properties of the cells, and the mutant phenotype is detectable only in the differentiated β-cells.

Our study clearly demonstrates that the mutation compromises the KCNQ1/Kv7 channel function. The mutated amino acid (R397W) of the KCNQ1 protein disrupts the secondary structure of Helix A. The C-terminal structures of Helix A and Helix B form a clamshell-like structure that is essential for calmodulin (CaM) binding, and this regulates the formation of the functional KCNQ1/Kv7 channel (Sun & MacKinnon, 2017). As a result, the mutation disrupts the generation of delayed outward K^+^ current (outward rectification) (Rorsman & Ashcroft, 2018), which in turn is predicted to reduce voltage-gated activation. This was confirmed in our Patch-clamp studies and *in vitro* electrophysiological analyses. The dysfunctional KCNQ1/Kv7 channel exhibits altered ATP sensitivity (Li, Gao et al., 2013), and this contributes to downstream processes. One way is by upregulating oxidative phosphorylation, which could modulate the activities of additional channels (e.g. KATP channels and CaV) in KCNQ1^R397W^ β-like cells. The altered Ca^2+^ influx leads to elevated levels of cytoplasmic Ca^2+^. As Ca^2+^ and insulin generate hexameric structures (Rorsman & Ashcroft, 2018, Dunn, 2005), this process promotes the crystallization of insulin into secretory insulin granules. Accumulated cytoplasmic Ca^2+^ thus triggers a cascade of insulin secretion through granule exocytosis in β cells (Fridlyand, Jacobson et al., 2013).

Over the long run, however, the above process decreases cellular metabolism, and ultimately causes cells to reduce their secretion of insulin. We also noted that a chronic exposure of KCNQ1^R397W^ β-like cells to high glucose promotes an irreversible deterioration of the cells in a way characteristic for diabetic conditions (Robertson et al., 2003). Overstimulation of cytosolic Ca^2+^ likely activates an additional downstream process, the NLPR3 inflammasome pathway (Murakami, Ockinger et al., 2012, Rossol, Pierer et al., 2012), that was reported to induce apoptosis *in vivo* (Bernal-Mizrachi et al., 2010, Kato et al., 2008, Epstein et al., 1989). Our comprehensive analysis of the KCNQ1^R397W^ organoid let us hypothesize that by birth, the impaired function of β-cells and the loss of β cell mass might have decreased the secretion of insulin into the patient’s blood to levels that were below detection. Thus, the homozygous C1189T mutation, identified in the patient might explain the phenotype of permanent neonatal diabetes (PND).

Several mutations of *KCNQ1*, which affect the C-terminal A and B Helices of KCNQ1, have been linked to irregular heartbeat (LQT1) (Ghosh et al., 2006), and the function of KCNQ1 as a voltage-gated potassium channel is well characterized for the repolarization phase of the cardiac action potential (Bellocq et al., 2004). In contrast, the exact role of KCNQ1 in glucose metabolism is still enigmatic. In a genetic survey for LQT1 syndrome, the 1189 C>T nucleotide change was classified as a mutation whose pathological significance was unknown (Kapplinger, Tester et al., 2009). However, the same mutation was reported as a loss of function and the potential cause of an intrauterine fetal death case (16-weak-old female), supported by a significant reduction (>70%) in IKs density in HEK273 cells using Patch-clamp (Crotti et al., 2013). This reduction is consistent with the *in vitro* electrophysiological phenotype of a mutation that causes LQT1 and impairments of channel functions in cardiomyocytes (Bellocq, van Ginneken et al., 2004). Here, our organoid model demonstrates clearly that this mutation of *KCNQ1* affects the rhythm of membrane depolarization/repolarization of β-cells and, as a result, their secretion of insulin. Furthermore, our patient’s condition of is stable under regular insulin treatment, and shows no sign of the cardiovascular phenotype. This suggests that the disease is caused by defects in the production of insulin and/or its secretion by β cells.

In fact, there is no clear connection between the cardiovascular and metabolic phenotypes associated with KCNQ1. The LQT1-syndrome has incomplete clinical penetrance in single families that carry heterozygous *KCNQ1* mutations. One explanation for the lack of clarity might be that the cardiovascular and metabolic syndromes appear at different stages of a patient’s life. In line with this, a study that monitored fourteen LQT1 patients with dominant-negative mutations of *KCNQ1* found that all developed postprandial hyper-insulinemia, but only at ages over 40 (Torekov, Iepsen et al., 2014). Our patient is younger than the onset of LQT1 syndrome (transitional period is 12 to 14 years old; fatal period is the median age of 32 (Guettler, Rajappan et al., 2019), which mean that it might simply be too early to draw any conclusions about the cardiovascular aspects of his disease.

Deciphering the exact role of KCNQ1/Kv7 channel in glucose metabolism was also hampered by the occasional contradictory reports on hypo- and hyper-insulinemic phenotypes. For example, it has been reported that knocking out KCNQ1 led to impaired GSIS in differentiated β-like cells in both mice and humans (Zeng et al., 2016, Boini et al., 2009). In contrast, blocking the KCNQ1/Kv7 channel and the following increase of cytoplasmic Ca^2+^ concentration results in hyper-insulinemia in rats (Finol-Urdaneta, Remedi et al., 2012). Furthermore, the forced expression of *KCNQ1* has been also associated with dysfunctional insulin exocytosis (Yamagata, Senokuchi et al., 2011). Importantly, our organoids could model both the hypo- and hyper-insulinemic phenotypes of the compromised KCNQ1. We could link these two phenotypes to the different maturation stages of β cells.

Species-specific differences in glucose metabolism may have contributed to the complexity of deciphering KCNQ1/Kv7’s roles in glucose metabolism; they may also explain some of the contradictions. Our animal disease model revealed that the impaired secretion of insulin exhibits only the hypo- (but not the hyper-) insulinemic phenotype in mouse β cells, in contrast to situation in humans. The decrease in insulin secretion may be due to a reduction in levels of expression of insulin, which is not observed in human β cells, suggesting a species-specific difference in the regulation of insulin production. We also detected a difference in the regulation of GLUT1/GLUT2 in human and mice β cells, which supports previous observations that GLUT1 is the primary glucose transporter in human pancreatic cells (De Vos et al., 1995). In mice β cells, both *Fgfrl1* and *Pdx1* have been shown to regulate the expression of *Glut2* (Hart et al., 2000, Waeber et al., 1996, Ahlgren et al., 1998, Brissova et al., 2002. In human β-like cells, by contrast, KCNQ1^R397W^ did not affect *GLUT2*. Instead, reductions of *FGFR1* and *PDX1,* downregulated *GLUT1* and directly impaired GSIS.

Finally, our study highlights that the pathogenic phenotype is a contingent one. While the same R397W mutation was reported from an intrauterine death case (Crotti et al., 2013), our patient survived, and so far, his heterozygotic family members are asymptomatic. We observed a downregulation of *HNF4α*, a MODY factor (Sanyoura et al., 2018), which might play a role in both the appearance and severity of the diseased phenotype. Potentially our study might help to decipher a potential link between mono- and polygenetic forms of diabetes (*KCNQ1*-PND, *HNF4α*-MODY and T2D). Additional factors, in conjunction with the homozygous KCNQ1^R397W^ mutation might accelerate the mass deterioration of β-cells, through (epi)genetic or environmental mechanisms that remain to be identified.

## Experimental Procedures

### Patients and genetic discovering

The patients were recruited from Charité. We excluded mutations in known genes causing neonatal diabetes (*ABCC8, KCNJ11, INS*) or hyperinsulinemic hypoglycemia (*GCK, HNF4A, GLUD1, ABCC8, KCNJ11*) by Sanger sequencing. We performed exome sequencing in the child with neonatal diabetes and his consanguineous parents using the Agilent SureSelect Human All Exon Kit (Agilent SureSelect v4, 50Mb). We analyzed the data with our established pipeline in the institute (Kuhnen, Turan et al., 2014) and confirmed the *KCNQ1* mutations by Sanger sequencing (Table S1) of the families. The Charité committee on human research approved the study (EA-No EA2/054/11) and written informed consent was obtained.

### Electrophysiology analysis of KCNQ1-R397W in CHO cells

Chinese hamster ovary (CHO) cells were cultured in Hams F12 medium supplemented with 10% FBS and 1% Strep/Pen in incubator. Cells were treated with trypsin-EDTA and plated on glass coverslips (Menzel, precoated with gelatin for 1h) in 6 well plates. We performed mutagenesis with the primers (Table S1) designed for the R397W mutation according to the manufacturer’s instruction (QuikChange II Site-Directed Mutagenesis Kit, Agilent Technologies, USA). Upon reaching 50-60% density, cells were transiently transfected with 2 µg human *KCNQ1* cDNA or 2 µg of human KCNQ1-R397W cDNA (pIRES-AcGFP1 vector, 19 cells) respectively, using SuperFect Transfection Reagent (Qiagen) according to the manufacture’s protocol. GFP protein expression was used to visually identify transfected CHO cells expressing KCNQ1 or KCNQ1-R397W. Patch-clamp recordings were obtained 48-72h after transfection.

Patch-clamp recorded (voltage-clamp whole-cell mode) at RT. The coverslips containing adherent transiently transfected CHO cells were transferred to a glass dish containing extracellular bath solution (135 mM NaCl, 5 mM KCl, 1.3 mM MgCl2, 5 mM HEPES, 2.5 mM CaCl2, 10 mM d-glucose, adjusted to pH 7.4 using NaOH). Pipettes were pulled from a vertical puller (Narishige, Tokyo) and maintained a resistance of 3-6 MΩ when filled with the intracellular solution (10 mM NaCl, 120 mM KCl, 2 mM MgCl2, 11 mM HEPES, 1 mM CaCl2, 11 mM EGTA, adjusted to pH 7.2 using KOH). The glass dish was placed under an inverted microscope equipped with fluorescence optics for green fluorescent protein detection (Olympus IX71 inverted microscope) to identify KCNQ1 and KCNQ1-R397W transfected CHO cells. Electrophysiological data were acquired via a Multiclamp 700B amplifier and a Digidata 1440A acquisition system. Test currents were elicited from a holding voltage of -100mV by potentials ranging from −100 mV to +80 mV in 20mV increments for KCNQ1 and the corresponding mutant KCNQ1-R397W. Leak subtraction was not applied. The individual cell current amplitudes were normalized to cell size (whole-cell membrane capacitance). Data were analyzed using pClamp 10.3 software (Molecular Devices) and statistical analysis (t-test, Mean ± SEM).

### Generation of KCNQ1^R397W^ mutant hESC cell lines

For an easy identification of the engineered cells, we used a CRISPR-Cas9 dependent homology-directed genome editing system that co-expresses a fluorescent marker (Richardson, Ray et al., 2016). The gRNA targeting the mutation of *KCNQ1* was designed and inserted into Px458-GFP cloning backbones (Addgene) for co-expression with Cas9. Single-stranded DNA (ssDNA) donors were designed as per Christopher’ protocol (Richardson et al., 2016). gRNA and ssDNA sequences used are listed in Table S2. The plasmids and ssDNA were transfected into hESC_H1 (WiCell) of 70-85% confluency using the XtremeGENE 9 transfection reagent (Roche). After 48 hours, the cells were dissociated to single-cell suspension using Accutase (StemPro) and sort GFP positive cells. To generate stable hESCs colonies from the fluorescent cell library, single colonies were picked and transferred to and cultured on Matrigel (Corning)-coated 48-well plates. To verify their genotype, we collected dead cells/debris floating in the culture media and PCR-amplified the CRISPR-edited genomic region. Amplicons were subjected to Sanger sequencing. Primers are listed in Table S3. Positive colonies and some unmodified colonies were continually cultured. Cell colonies were separately dissociated and transferred into 6-well plates using ReLeSR (Stem Cell Technologies). Differentiated cells were removed by using a pipette tip to minimize spontaneous differentiation. The high pluripotent stem cells form colonies were selected by Anti-Tra-1-60-PE and Anti-PE MicroBeads following the manufacturer’s instructions (MACS Miltenyi Biotec). The stable colonies were frozen down at -80°C. To eliminate the potential off-target events generated by the CRISPR system, we established one unmodified control (UC) and generated two *KCNQ1* mutant clones (KCNQ1^R397W^ No.1 and No.2) from the CRISPR/Cas9 transfected cell library. Robert Koch institute approved the associated studies in hESC_H1 (AZ: 3.04.02/0147).

### Generation of human pancreatic β-like cells

hESCs were developed toward insulin+ cells in a suspension-based format on a shaker with minor modification to the published protocol (Chiou et al., 2021, Velazco-Cruz et al., 2019). The single cells were seeded in mTeSR1 media (Stem Cell Technologies) supplemented with CloneR (Stem Cell Technologies) in 6-well ultra-low attachment plates at 5.5 × 10^6^ cells/well. The plates were cultured on the shaker (Binder) at 100 rpm in a CO2 incubator (Binder) for 24 h. Undifferentiated aggregates were cultured in daily differentiation media.

S1/S2 basal media: 500 mL MCDB131 (Life Technologies) supplemented with 0.75 g NaHCO3, 1% GlutaMAX (Life Technologies), 15 mM glucose (Sigma) and 2.5 g fatty acid-free BSA (Proliant Biologicals).

S3/S4 basal media: 500 mL MCDB131 supplemented with 1.25 g NaHCO3, 1% GlutaMAX, 15 mM glucose and 10 g fatty acid-free BSA.

S5/S6 basal media: 500 mL MCDB131 supplemented with 0.75 g NaHCO3, 1% GlutaMAX, 20 mM glucose and 10 g fatty acid-free BSA.

Day 0 media: S1/S2 basal media, 100 ng/mL Activin A (R&D Systems), 25 ng/mL mouse Wnt3a (R&D Systems).

Day 1 - Day 2 media: S1/S2 basal media, 100 ng/mL Activin A.

Day 3 - Day 5 media: S1/S2 basal media, 50 ng/mL KGF (R&D Systems), 0.25 mM ascorbic acid (Sigma).

Day 6 - Day 7 media: S3/S4 basal media, 50ng/mL KGF, 0.25 μM SANT-1 (Sigma), 1 μM RA (Sigma), 100 nM LDN-193189 (Stemgent), 200 nM TPB (EMD Millipore), 0.25 mM ascorbic acid, 0.5% ITS-X (ThermoFisher).

The plates were cultured on the shaker at 120 rpm in CO2 incubator from day 8 to day 20.

Day 8 - Day 10 media: S3/S4 basal media, 2ng/mL KGF, 0.25 μM SANT-1, 0.1 μM RA, 200 nM LDN-193189, 100 nM TPB, 0.25 mM ascorbic acid, 0.5% ITS-X.

Day 11 - Day 13 media: S5/S6 basal media, 0.25 μM SANT-1, 0.05 μM RA, 100 nM LDN-193189, 1 μM T3 (Sigma), 10 μM ALK5i II (Enzo Life Sciences), 10 μM ZnSO4 (Sigma), 10 μg/mL heparin (Sigma), 0.25 mM ascorbic acid, 0.5% ITS-X.

Day 14 - Day 20 media: S5/S6 basal media, 100 nM LDN-193189, 1 μM T3, 10 μM ALK5i II, 10 μM ZnSO4, 10 μg/mL heparin, 100nM ɣ-secretase inhibitor XX (Calbiochem), 0.5% ITS-X.

S7 media (day 21-day 40): 500 mL MCDB131 supplemented with 1% GlutaMAX, 10 g fatty acid-free BSA, 5mg heparin, 5mL MEM nonessential amino acids, 84 μg ZnSO4, 500 μL Trace Elements A and 500 μL Trace Elements B. Day 21 aggregates were dissociated to single cells and were seeded in S7 media supplemented with CloneR in 6-well ultra-low attachment plates at 5.5 × 10^6^ cells/well. The plates were cultured on the shaker at 100 rpm in a CO2 incubator for 24 h. Day22 aggregates were cultured in standard S7 media without CloneR at 120 rpm.

### Flow cytometry analysis

The cell aggregates were treated with TrypLE (10X, ThermoFisher) to dissociate to single cells. The single cells were re-suspended with cold BD fixation/permeabilization solution following the manufacturer’s instructions (BD Biosciences). Quality control of the differentiation was performed by flow cytometry analysis of stage specific markers (Table S4). Corresponding isotypes antibodies were loaded into another suspension aliquot as isotope control. Cells were washed and suspended into 0.2% BSA after aspirating supernatant. The samples were analyzed on a flow cytometer after the compensation setting.

### Immunofluorescence analysis

Organoids were fixed in 4% PFA and dehydrated in 30% sucrose (w/v). Organoids were transferred to the center of flat bottom cryosectioning molds (VWR). The mold was filled with OCT and placed in a dry ice ethanol bath to freeze OCT (VWR). Organoids were embedded in frozen OCT and stored at -80°C. The embedded organoids were sectioned by using CRYOSTAT MICROTOM (Thermo Scientific).

Sectioned slides were washed by DPBS to get rid of OCT. The slide was incubated with respective primary antibody and second antibody solutions after blocking. The nuclei were stained with DAPI (fisher scientific). The slides were mounted with VECTASHIELD® Antifade Mounting Medium (Vector Laboratories) and covered with coverslips. The mounted and covered slide was sealed with CoverGripTM Coverslip Sealant (Biotium) and allowed to dry fully before analyzed on LSM700 inverted fluorescent microscope (Zeiss). Depending on the antibody combinations, the 405, 488, 555 and 640 nm excitation lasers were used in sequential scans to prevent cross-talk between the detection fluorescence channels. The slides were stored long-term at -80°C. Antibodies used are listed in Table S4.

### Gene expression analysis (qRT-PCR)

Total RNA was extracted from cells using the Direct-zol RNA MiniPrep Plus kit following the manufacturer’s instructions (Zymo research) and used for cDNA reverse transcription (Applied Biosystems). Gene expression was assessed on the 7900HT Fast Real-Time PCR System (Applied Biosystems) using the Power SYBR Green PCR Master Mix (Applied Biosystems). Data were normalized to *GAPDH* or *TBP* expression using the △△Ct method. Primers used are listed in Table S3.

### *KCNQ1* mutation locus methylation analysis

The cell aggregates were lysed in lysis buffer (100 mM Tirs-HCl, 0.5 M EDTA, 10%SDS, 5 M NaCl, 0.05% Protein K) and incubated at 55°C overnight. The lysate was mixed with an equal volume of phenol: chloroform: isoamyl alcohol solution (Roch). The aqueous phase (upper) was mixed with a 10% volume of 3 M sodium acetate (pH 5.2) and a 2-fold volume of -20°C cold 100% ethanol. The mixture was placed at -80°C overnight. The supernatant was carefully removed after DNA was settled down by gravity. The DNA pellet was washed with 70% ethanol. The DNA pellet was allowed to air dry for 15 min before resuspending in Nuclear free H2O (Sigma). Sodium bisulfite conversion of unmethylated cytosines in DNA was based on EpiTect Bisulfite Handbook (Qiagen). We designed primers (Table S3) from MethPrimer to sequence the *KCNQ1* mutation locus.

### Western blotting

Organoids were lysed in RIPA buffer (50 mM Tris-HCl pH7.4, 150 mM NaCl, 1 mM EDTA, 1% Triton-100, 1% Na-Deoxycholate and 0.1% SDS). The procedure of protein concentration determination was based on the manual of the Pierce BCA Protein Assay Kit (Pierce). Protein samples were boiled at 95°C for 5 min and were run on a TGX Stain-Free acrylamide gel. The gel was prepared by following the manual of the TGX Stain-Free FastCast Acrylamide Kit (BioRad). The proteins were transferred onto a PVDF membrane (BioRad) following the guide of the Trans-Blot Turbo transfer system RTA Transfer Kit (BioRad). The PVDF membrane was blocked and then incubated with KCNQ1 antibodies (ATLAS ANTIBODIES) overnight at 4°C. The PVDF membrane was washed with TBST buffer and incubated with HPR-Anti rabbit lgG (Thermo Scientific) for 1h at room temperature. For detection of KCNQ1, the PVDF membrane was developed with SuperSignal West Femto Maximum Sensitivity Substrate (Thermo Scientific). The antibodies were removed by the mild stripping buffer (15 g Glycine, 1 g SDS, 10 mL Tween 20, add ddH2O to 1 L, pH 2.2). The PVDF membrane was blocked and incubated with Actin antibodies (Dianova) overnight at 4°C. The PVDF membrane was incubated with HPR-Anti mouse lgG (Thermo Scientific) for 1h at room temperature. For detection of Actin, the membrane was developed with ECL reagents (Cytiva). The PVDF membrane was imaged on the ChemiDocTM MP imaging system (BioRad).

### Insulin secretion analysis of human KCNQ1^R397W^ β-like cells

The organoids were transferred to 6-well ultra-low attachment plates with 5 mL KRB buffer (130 mM NaCl, 5 mM KCl, 1.2 mM CaCl2, 1.2 mM MgCl2,1.2 mM KH2PO4, 20 mM Hepes (pH 7.4), 25 mM NaHCO3, 0.1% BSA) containing 2.75 mM glucose. The plates were incubated for 1 h in a 37℃ incubator. 5 organoids/well were transferred to 96-well plates with 10 replicates.

Insulin secretion stimulated by glucose or KCl was firstly measured by adding 2.75 mM glucose KRB buffer until 200 μL/well. The organoids were incubated 30 min in a 37℃ incubator. 190 μL solution per well was transferred into a new 96-well plate, respectively. Fresh 190 μL 16.8 mM glucose KRB buffer solution or 30 mM KCl KRB buffer per well was added back to the previous well following by incubation for 30 min in a 37℃ incubator. 190 μL solution per well was transferred into a new well of 96-well plate, respectively. Total insulin was measured by adding the acid ethanol solution (1.5% HCl, 80% ethanol) until 200 μL/well. The organoids were incubated at 4℃ overnight on the shaker at 100 rpm. 190 μL solution per well was transferred into a new well of 96-well plate, respectively. The new 96-well plates were frozen at -80°C for ELISA measurement.

The released and total insulin was measure by the Human insulin ELISA Kit following the manufacturer’s instructions (ALPCO). The organoids were collected into 50 μL/well sonication buffer (10 mM Tirs, 1 mM EDTA, 0.2% Triton-X 100, 0.05% Protein K). The organoids were sonicated 5 cycles (30 s ON and 30 s OFF) in Bioruptor Pico Sonication device (diagenode). The DNA of organoids was detected by using dsDNA Broad Range Assay (DeNovix). The DNA concentration was calculated by using DeNovix DS-11 FX Fluorometer (DeNovix). Insulin secretion was normalized to low glucose stimulation, and insulin content was normalized to DNA mass.

### Insulin secretion analysis of KCNQ1^R397W^ transfected mouse pancreatic β cells

The hKCNQ1 cDNA was subcloned into the pIRES2-AcGFP1 vector. We transfected the *KCNQ1* constructs and the empty vector into our established mouse pancreatic β cells (referred to as SJ β cells (Jia et al., 2015)) using AmaxaTM Cell Line NucleofectorTM Kit V and *AmaxaTM* NucleofectorTM 2b device (Lonza, Switzerland). The glucose-sensitive mouse β-cell line SJ-β was cultured and an insulin-secretion assay performed as reported earlier (Jia et al., 2015). For control, cells were transfected with wild-type *KCNQ1* or left untransfected. Insulin secreted into medium and from whole-cell lysates was quantified with Mouse High Range Insulin ELISA kit, according to manufacturer’s protocol (Alpco, Salem, USA). Cells were incubated for 30 min with low (3.3mM) or high (16.7mM) glucose concentration, and 30mM of KCl to depolarize the cell membrane. We normalized secreted insulin by the total cellular insulin and total insulin was normalized by the total cell protein. The experiments were repeated four times with different patches of mouse β-cells.

### Electrical activity analysis

Organoids were treated with TrypLE (10X) to dissociate to single cells. The 5% Matrigel solution was prepared by diluting Matrigel in S7 media supplemented with CloneR. The cell pellet was re-suspended with 10 μL cold 5% Matrigel after aspirating supernatant. 8 μL cell suspension was dotted to recording electrodes avoiding ground electrodes and incubated 0.5 h in a CO2 incubator until cell attachment. 300 μL S7 media supplemented with CloneR was added against the wall of the well. The electrical activity (Spike Detector) was recorded in Neural Spikes mode on MAESTRO Pro (AXION BIOSYSTEMS). The threshold-baseline of Spike Detector was set by using 3 mM glucose.

### Cytoplasmic Ca^2+^ level measurement

Human islet and hESC-derived differentiated organoids (approximately 20 organoids or human islets per well) were respectively plated into a 96-well black plate (ThermoFisher) coated with 1% hESC-qualified Matrigel. After 24 h, the wells were washed with prewarmed (37 ℃) KRB buffer containing 2.5 mM glucose. The cell organoids were incubated with 50 μM Ca^2+^-sensitive fluorescent probe Fluo4-AM (Life Techbologies) in 2.5 mM glucose KRB buffer for 45 min in a 37℃ incubator. The plate was incubated further in a 37℃ incubator for 15 min after washing with 2.5 mM glucose KRB buffer. The plate was immediately staged on a Cell R live Imaging System (Olympus) to acquire time-series imaging.

Fluo-4 AM was illuminated at using an excitation filter 492/18 nm, and its emission was collected between 500-550 nm. Time-series images were recorded at a resolution of 80x magnification with a 20-sec interval. The progression of glucose challenges and time of the stimulation during imaging was as follows: Imaging started after 5 min incubation in KRB buffer containing 2 mM glucose and ran 16 cycles. The next step was followed by a 5 min incubation in KRB buffer containing 20 mM glucose and ran 16 cycles. Sequential low and high glucose challenges were repeated one more time after washing with low glucose KRB buffer. The organoids’ imaging was resumed after adding low or high glucose solution. Fluorescence intensity was measured by using Fiji software. StackReg was applied to anchor the organoids throughout the imaging. The positions of organoids were added to the ROI manager.

The fluorescence intensity of the organoids was measured throughout the imaging. Finally, all of the fluorescence intensity of the same organoid was normalized to its first image.

### Electron Microscopy

β-cell like organoids (day 31) were fixed in a freshly prepared mixture of 2 % formaldehyde and 2 % glutaraldehyde (Sigma) in 0.1 M phosphate buffer (18.2% 0.1 M KH2PO4, 81.8% 0.1 M Na2HPO4 in ddH2O) for 1 h at room temperature, followed by fixation at 4°C overnight. Samples were stained with 1% OsO4 for 2 h after washing with 0.1 M phosphate buffer. They were dehydrated in a graded ethanol series and propylene oxide and embedded in Poly/Bed^R^ 812 (Polysciences Inc.). Ultrathin sections were contrasted with uranyl acetate and lead citrate. Finally, sections were examined with a Morgagni electron microscope (Thermo Fisher). Digital images were taken with a Morada CCD camera and the iTEM software (EMSIS GmbH, Münster).

### RNA-seq and data analysis

mRNA quality was checked by using Agilent 2100 Bioanalyzer following the protocol of RNA 6000 Nano Kit. BGI Hongkong prepared the DNA libraries and sequenced the libraries on a DNBseq Eukaryotic-T resequencing.

A 30 million 100 bp paired-end reads were obtained per sample. Discarding low-quality reads, trimming adaptor sequences, and eliminating poor-quality bases were done using FASTX-Toolkit and Trimmomatic. Building index and alignment, the reads were performed using Salmon after discarding outliers with over 30% disagreement. GC content and gene length biases were checked using R package NOISeq to quality control of count data. Mean-variance and PCA were calculated between biological replicates using the tximport package in R. The parameters of lengthscaledTPM were CPM cutoff >2 and sample cutoff 2 between the replicates for the analyzed groups. The RUv package from Bioconductor was used to eliminate batch effects. Therefore, all of the samples were normalized to TMM (weighted trimmed mean of M-values). Gene counts were used for differential expression analysis using the EdgeR package (Table S5). The gene ontology enrichment was performed using ShinyGo v0.61(KEGG, FDR 0.1). The common genes from categories were selected and made heatmaps using the complexheatmap package from Bioconductor (Gu, Eils et al., 2016).

### Pro-apoptosis analysis

Organoids were treated with TrypLE (10X) to dissociate to single cells and then washed with 1 mL 0.2% BSA. The cell pellet was re-suspended with 2 mL S7 media supplemented with CloneR. The cells were incubated in a 1% Matrigel coated plate for two days in a 37℃ incubator. The flat culture cells were treated with TrypLE (10X) to dissociate to single cells and then washed with 1 mL 0.2% BSA. FITC Annexin V antibodies and 7-AAD were loaded into 100 μL cells suspension aliquot to analyze pro-apoptosis following the manufacturer’s instructions (BD Biosciences). FITC isotypes were used as the isotope control. Cells were suspended into 200 μL 1X binding buffer after aspirating supernatant. One well of cells were treated with 200 μM H2O2 6 h as a positive apoptosis sample for compensation setting. The samples were analyzed on a flow cytometer after the compensation setting.

### Statistical analysis

All qRT-PCR data were analyzed by the △△Ct method. Data were analysed for normal distribution where applicable. Data were analyzed using unpaired/paired t-tests and one/two-way ANOVA. Dynamic Ca^2+^ flux was analyzed using two-way ANOVA and was generated with Prism8. The DNA sequence was visualized on Benchling. FlowJo was used to analyze Flow cytometry data. Immunofluorescence analysis was performed with Fiji. The intensity of protein on the PVDF membrane was analyzed with GelAnalyzer.

### Accession numbers

RNA-seq data are available from the GEO database under accession numbers GSE168245.

## Acknowledgements

We thank Dr. Sebastian Diecke (MDC) for preliminary experiments in hiPSCs. We thank Prof. G. Abbott (University of California USA) providing the human *KCNQ1* cDNA. We are grateful to Russell Hodge (MDC) for providing comments on the manuscript. We thank BioRender support the software for figure generation. We are also grateful to MDC for providing training grants. The staffs in the MDC facility are acknowledged for general maintenance and support. This work was supported by National Institutes of Health grants R01DK068471 and UG3DK122639 to M.S.

## Author Contributions

Zhimin Zhou designed and conducted the overall experiments and analyzed the data of hESCs-derived samples and human islets, and cowrote the manuscript. Maolian Gong contributed to the family genetic analysis, supervised insulin secretion analysis of mouse SJ-β cells. Amit Pande contributed to the RNA-seq data. Ulrike Lisewski and Torsten Röpke contributed to electrophysiology analysis in CHO cells. Bettina Purfürst contributed to the generation of electron microscopy images. Lei Liang and Shiqi Jia contributed insulin secretion analysis of mouse SJ-β cells. Sebastian Froehler and Wei Chen contributed to exome sequencing. Anca Margineanu assisted in calcium signaling imaging. Chun Zeng and Han Zhu provided advice on experimental design and hESC differentiation. Peter Kühnen, Semik Khodaverdi and Winfried Krill contributed to the clinical studies. Maike Sander supervised hESC differentiation into beta cells and provided comments on the manuscript. Klemens Raile contribute to the clinical studies, funding acquisition and supervision. Zsuzsanna Izsvak supervised the study, funding acquisition and cowrote the manuscript. Z.I., M.G. and K.R. generated the project. C.Z, H.Z, M.G. and K.R. provided comments and edits for the manuscript. All of the authors discussed the results and commented on the manuscript.

---

**Figure EV1.**
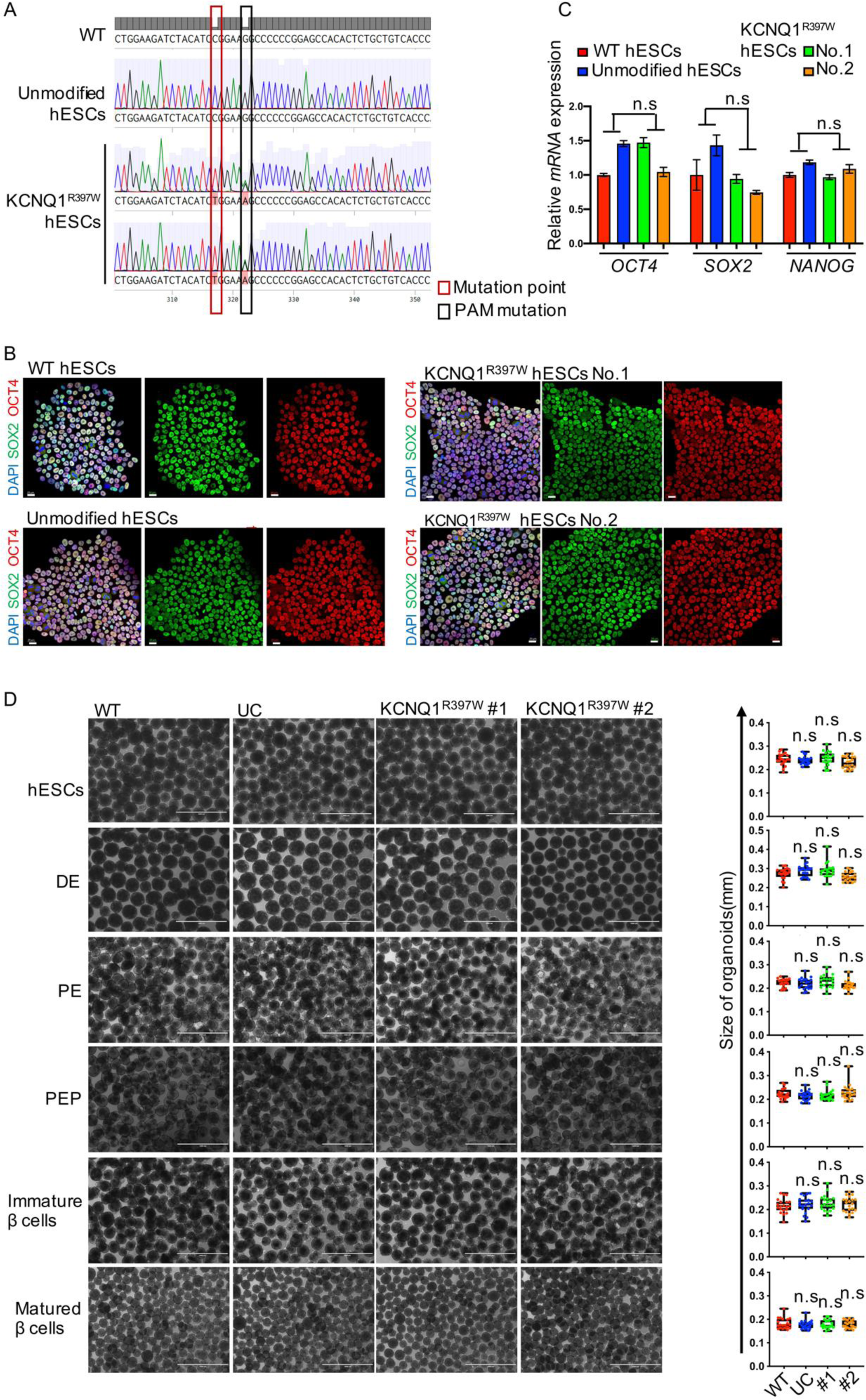
*In vitro* generation of the permanent neonatal diabetes (PND) patient*-*derived *KCNQ1* mutation (1189 C>T) in human β-like cells. A. Sanger sequencing of CRISPR/Cas9 edited KCNQ1^R397W^ mutation in hESCs (red box). PAM, protospacer adjacent motif (black box). B. Immunostaining of WT, UC and KCNQ1^R397W^ hESCs using pluripotency markers SOX2 and OCT4. Scale bar=20 μm. C. Characterization of WT, unmodified control (UC), and KCNQ1^R397W^ hESCs by qRT-PCR, specific for pluripotency markers (*OCT4*, *SOX2*, and *NANOG*). Data are normalized to *GAPDH* and presented as mean ± SD. D. Cell morphology at different stages of the differentiation, including the stage of hESCs (day 0), definitive endoderm (DE, day 3), pancreatic endoderm (PE, day 11), pancreatic endocrine precursors (PEP, day 14), immature β cells (day 21), and matured β cells (day 31). Scale bar=1 mm. Data information: In (C and D), data are presented as mean ± SD. n.s indicates a non-significant difference, *p < 0.05, **p < 0.01, ***p < 0.001, and ****p < 0.0001 (Student’s t-test).

**Figure EV2.**
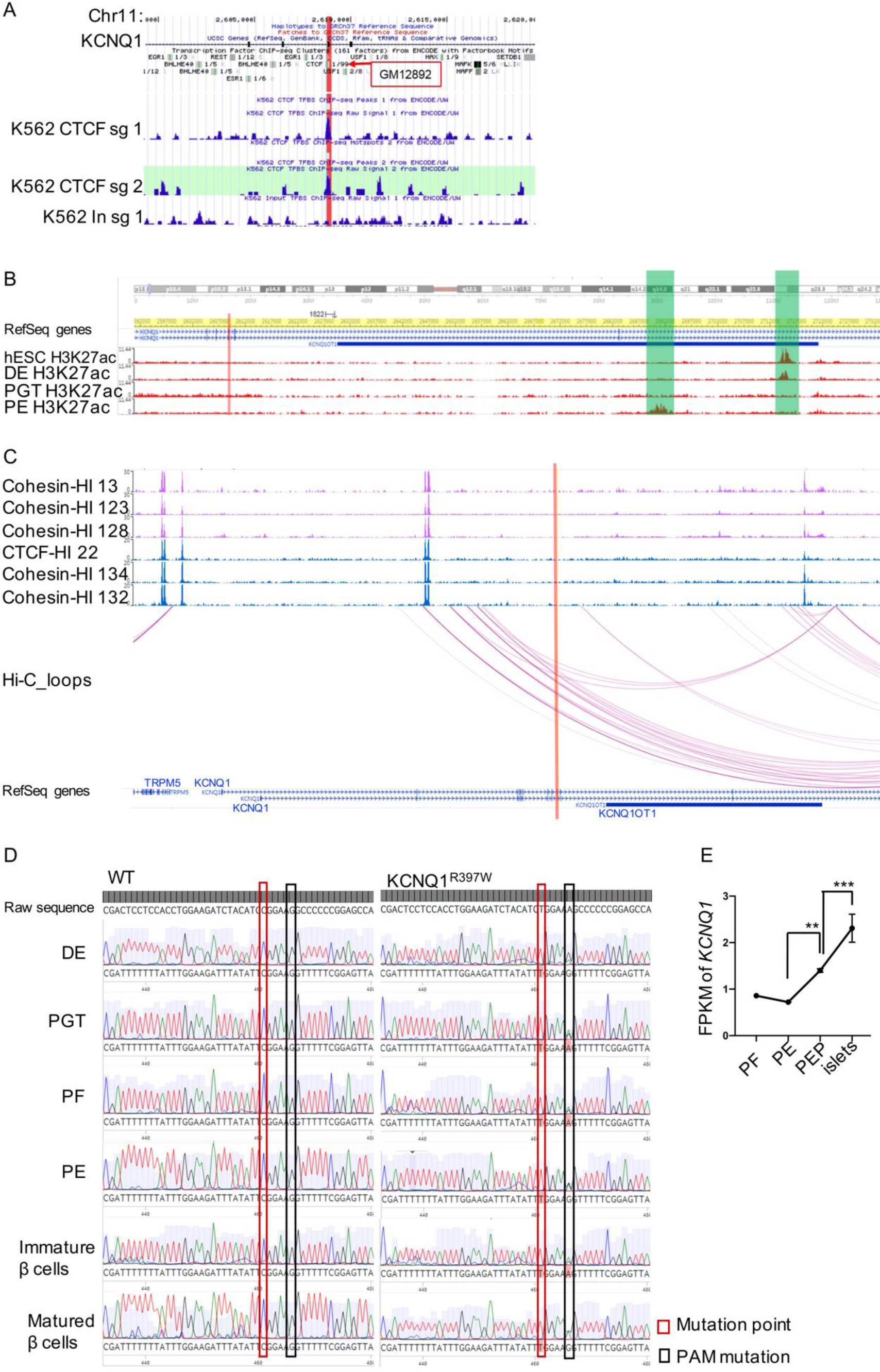
Regulatory genomics of the C1189T mutation of KCNQ1. A. The CTCF binding signal in KCNQ1 locus showed by UCSC Genome Browser. Tracks show signals of CTCF binding (potential CTCF binding motif) in cell lines: GM12892 (human B-lymphocyte, lymphoblastoid and K562 (human myelogenous leukemia). B. The dynamic enhancer signal during β-cell differentiation is revealed by H3K27ac CHIP assay(Xie et al., 2013) in hESCs, DE, primitive gut tube (PGT), and PE. The potential enhancer is highlighted in light green. C. Cohesin and CTCF binding (ChIP) signals and the predicted chromatin loops at the KCNQ1 gene and its neighboring genomic region in human islets (Hi-C data(Miguel-Escalada et al., 2019)). D. Cytosine methylation analysis in different stages of differentiation, including DE, PGT, posterior foregut (PF), PE, immature β cells and matured β cells. The C1189T mutation is boxed in red, whereas the PAM mutation marked by black. E. FPKM (Fragments per kilobase of transcript per million mapped reads) levels of *KCNQ1* transcripts during human pancreatic differentiation, in PF, PE, PEP and matured islets after transplanted into mice(Xie et al., 2013). Data presented as mean ± SD. n.s indicates a non-significant difference, *p < 0.05, **p < 0.01, ***p < 0.001, and ****p < 0.0001 (one-way ANOVA). Data information: In (A-C), the position of the C1189T mutation in exon 9 of KCNQ1 is marked by a red bar.

**Figure EV3.**
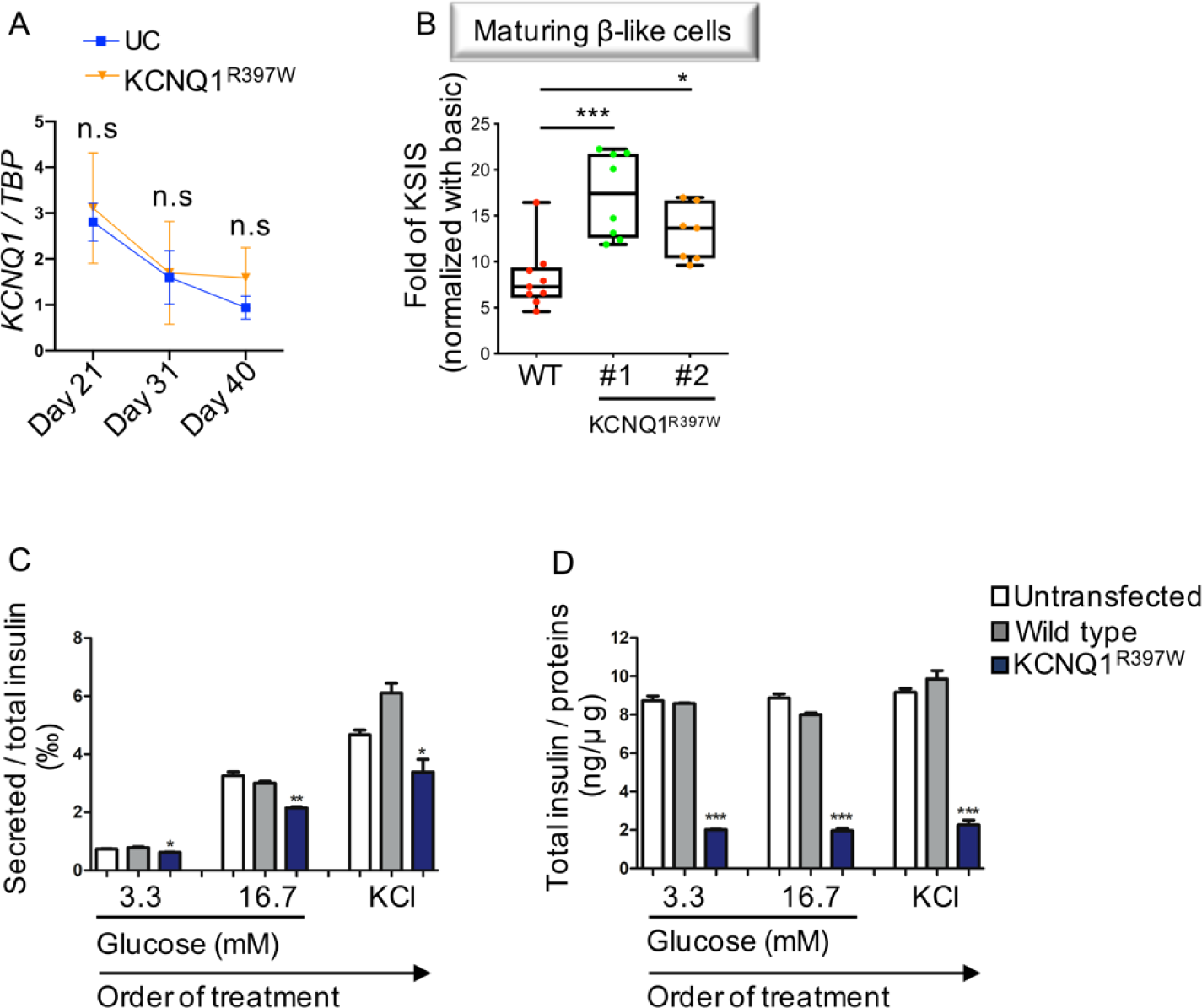
The insulin secretion of human day 28 organoids and KCNQ1^R397W^ transfected mouse SJ β cells upon stimulation. A. qRT-PCR analysis of the *KCNQ1* expression in human pancreatic organoids of UC, and KCNQ1^R397W^ β-like cells. p values calculated by two-way ANOVA. B. Fold-change of insulin secretion in maturing human β-like cells (day 28) with 30 mM KCl stimulation between WT and KCNQ1^R397W^ β-like cells. C, D. Insulin secretion assay (C), and total insulin assay (D) are analyzed with ELISA in mouse SJ β cells untransfected and transfected with expression constructs of KCNQ1^WT^ (wild type) and KCNQ1^R397W^. Data information: Data are presented as mean ± SD. n.s indicates a non-significant difference, *p < 0.05, **p < 0.01, ***p < 0.001, and ****p < 0.0001 (Student’s t-test).

**Figure EV4.**
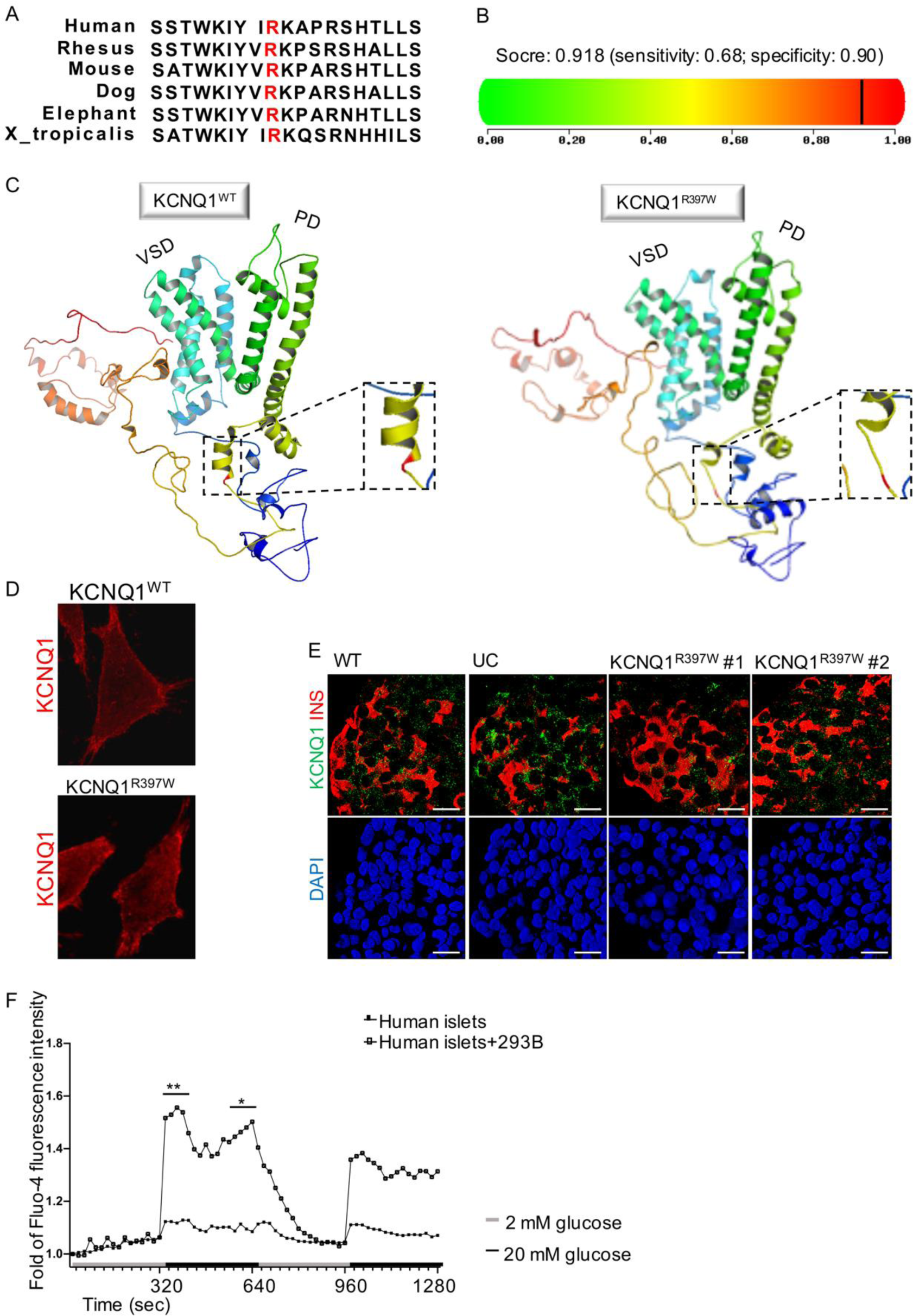
Function prediction of KCNQ1^R397W^. A. Amino acid sequence around the mutation (red) is conserved in KCNQ1. B. Polyphen2 prediction for the destructive potential of the R397W mutation in KCNQ1. C. 3D structure prediction of KCNQ1^WT^ and KCNQ1^R397W^. The figures are generated using the PyMOL viewer. Left panel, KCNQ1^WT^ with the mutation locus (highlighted in red) is located in an α helix. Right panel, KCNQ1^R397W^, the mutation disrupts the helical structure. Instead of an α helix, a random coil is predicted (The mutation is highlighted in red). D. Immunostaining for KCNQ1^WT^ and KCNQ1^R397W^ expressed in KCNQ1-null Chinese hamster ovary (CHO) cells (red). E. Immunostaining of insulin (red) and KCNQ1 (green) in human β-like cells (day 31) shown by 3D confocal scanning. Scale bar=20 μm. F. The dynamic Ca^2+^ flux of human β-cells by Fluo-4 AM staining. Human islets were cultured with or without Chromanol 293B (293B). Data are presented as mean. p values are calculated by two-way ANOVA. n.s indicates a non-significant difference, *p < 0.05, **p < 0.01, ***p < 0.001, and ****p < 0.0001.

**Figure EV5.**
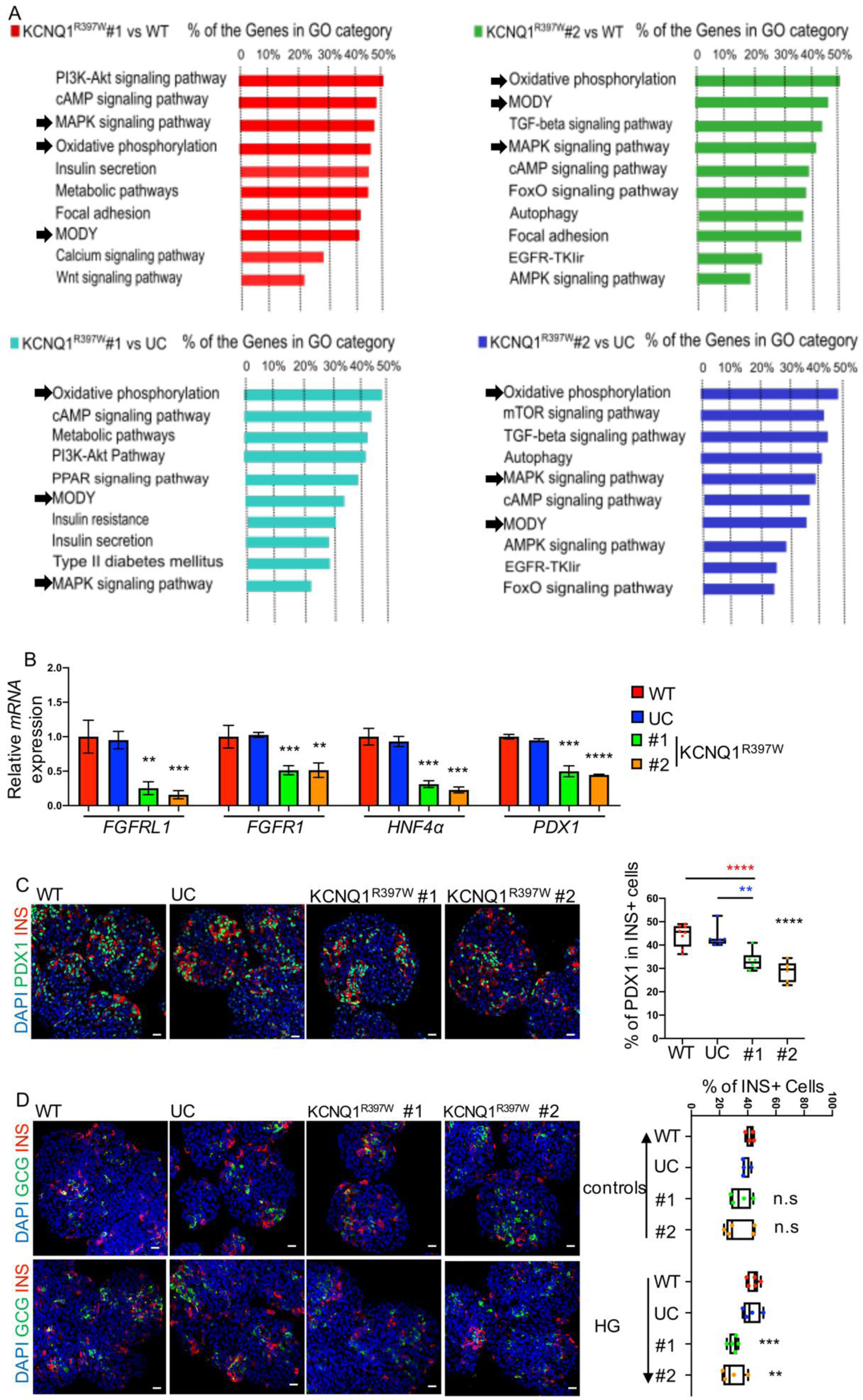
The analysis of differentially expressed genes, and the assay of chronic exposure to high glucose. A. GO analysis of differentially expressed genes (DEGs). The most significant categories are marked. B. qRT-PCR analysis of *FGFRL1*, *FGFR1*, *HNF4α* and *PDX1* expression in WT, UC, and KCNQ1^R397W^ β-like cells at their late stage (day 40). Data are normalized to *TBP*. C. Late-stage β-like cells were immunoassayed for PDX1 (green) and insulin (INS, red), and quantified. D. Late stage β-like cells were cultured in normal media (controls) or in media supplemented with 20 mM glucose (high glucose, HG), immunoassayed for glucagon (GCG, green) and insulin (INS, red), and quantified. Data information: Scale bar=20 μm (C and D). In (B-D), the data are presented as mean ± SD; n.s indicates a non-significant difference, *p < 0.05, **p < 0.01, ***p < 0.001, and ****p < 0.0001 (Student’s t-test).

**Table S1:**
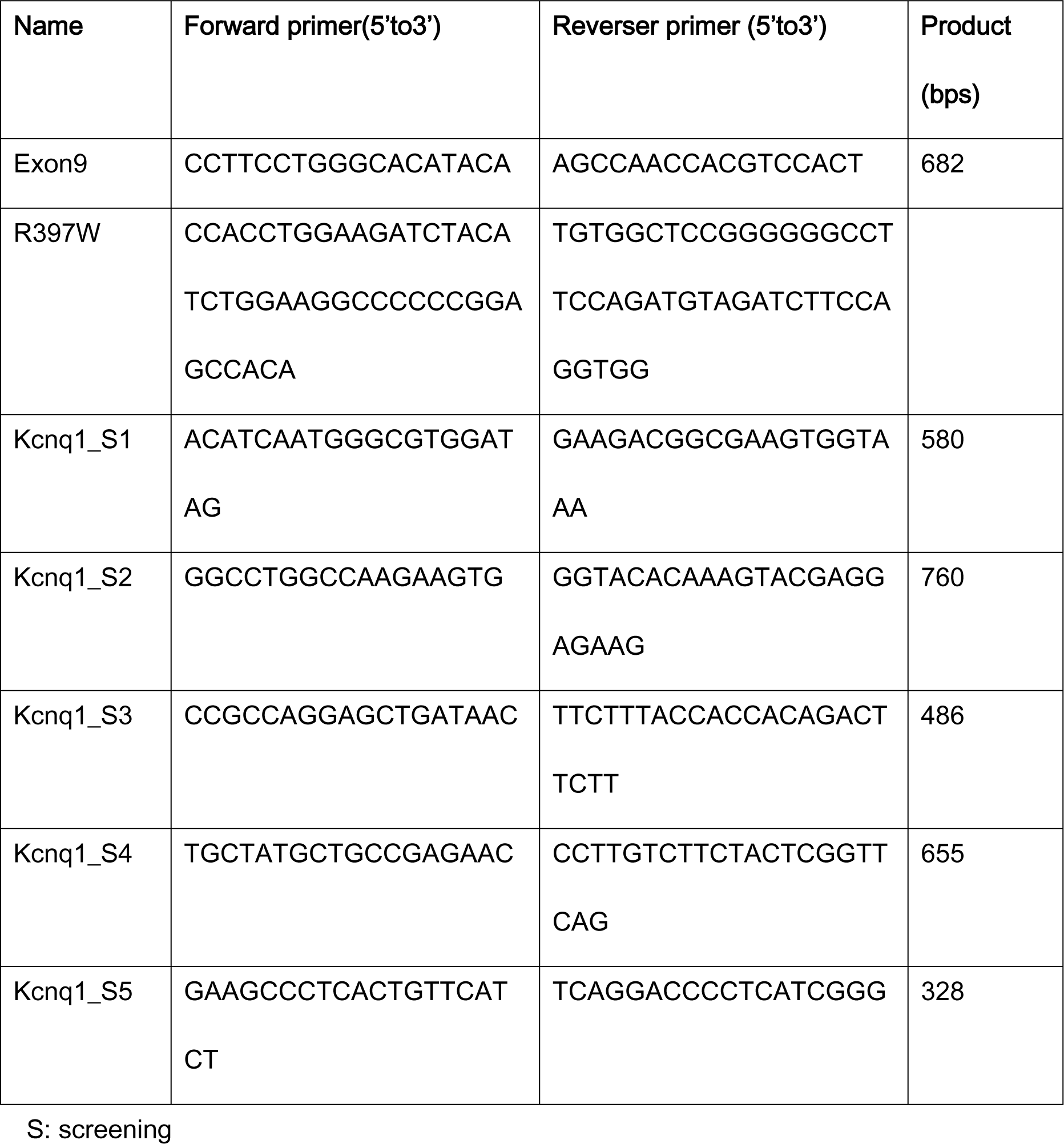
KCNQ1 primers for mutation confirmation, mutagenesis, and construct screening

**Table S2:**
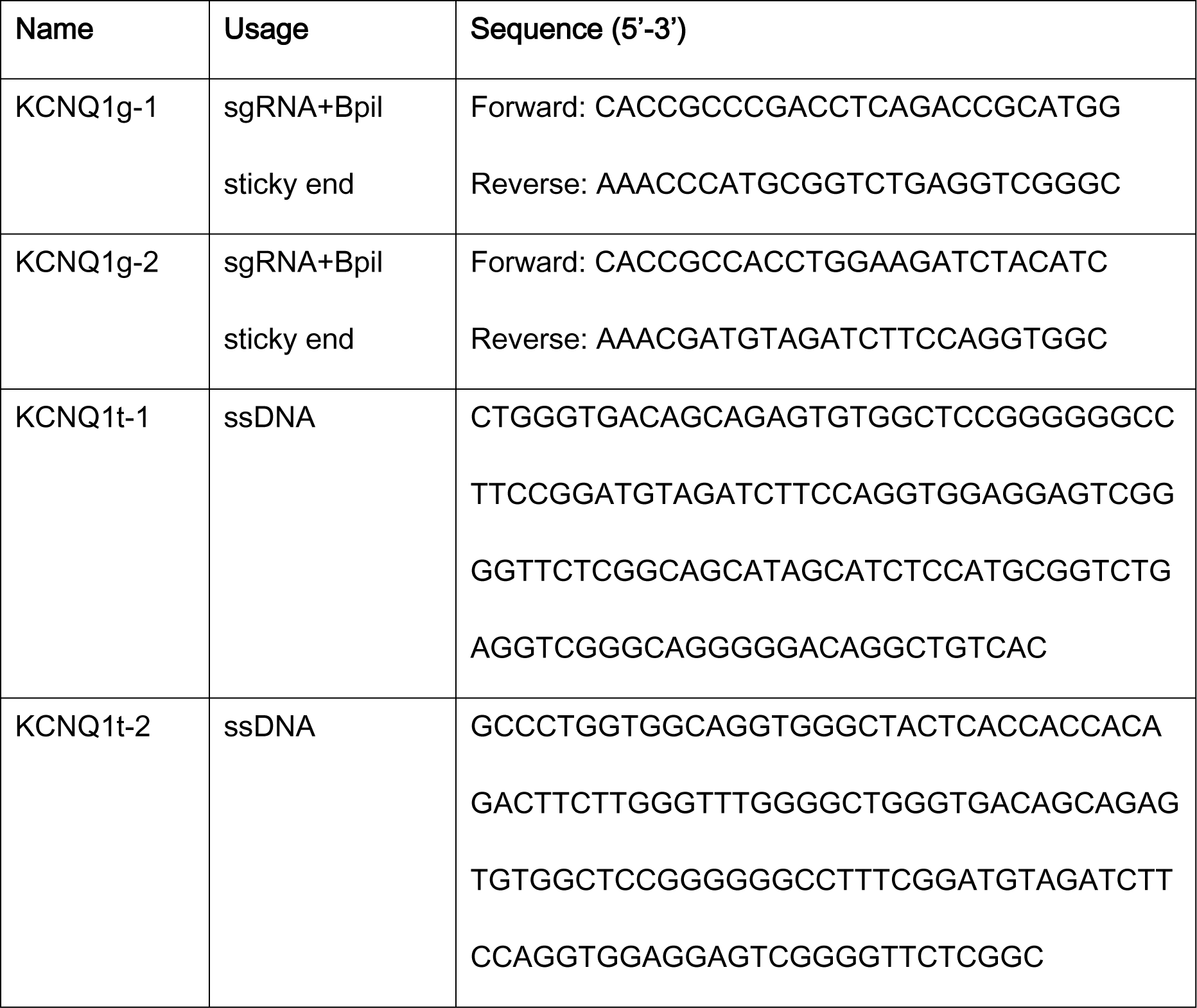
The sequence of gRNA and ssDNA for gene editing.

**Table S3:**
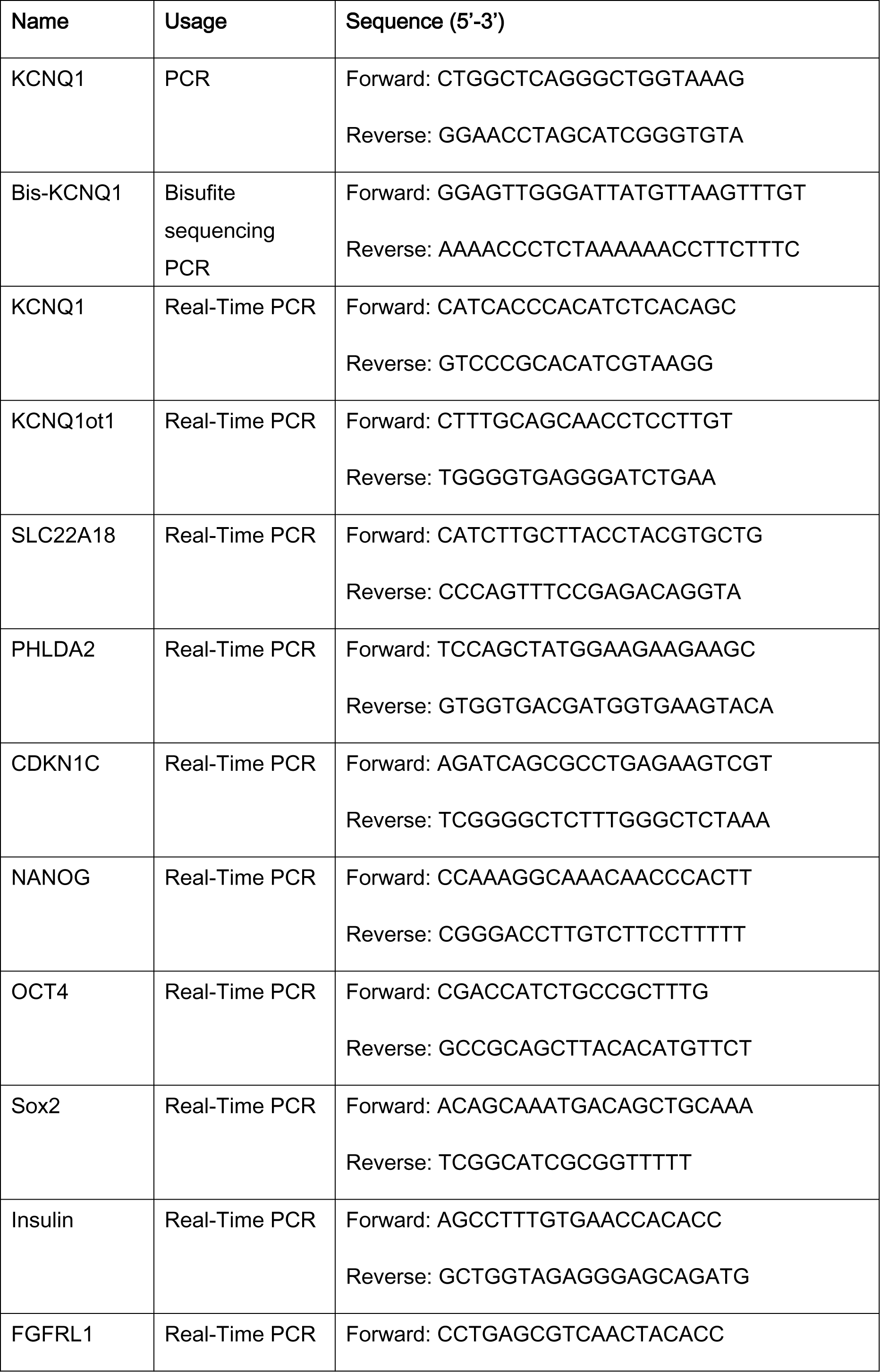
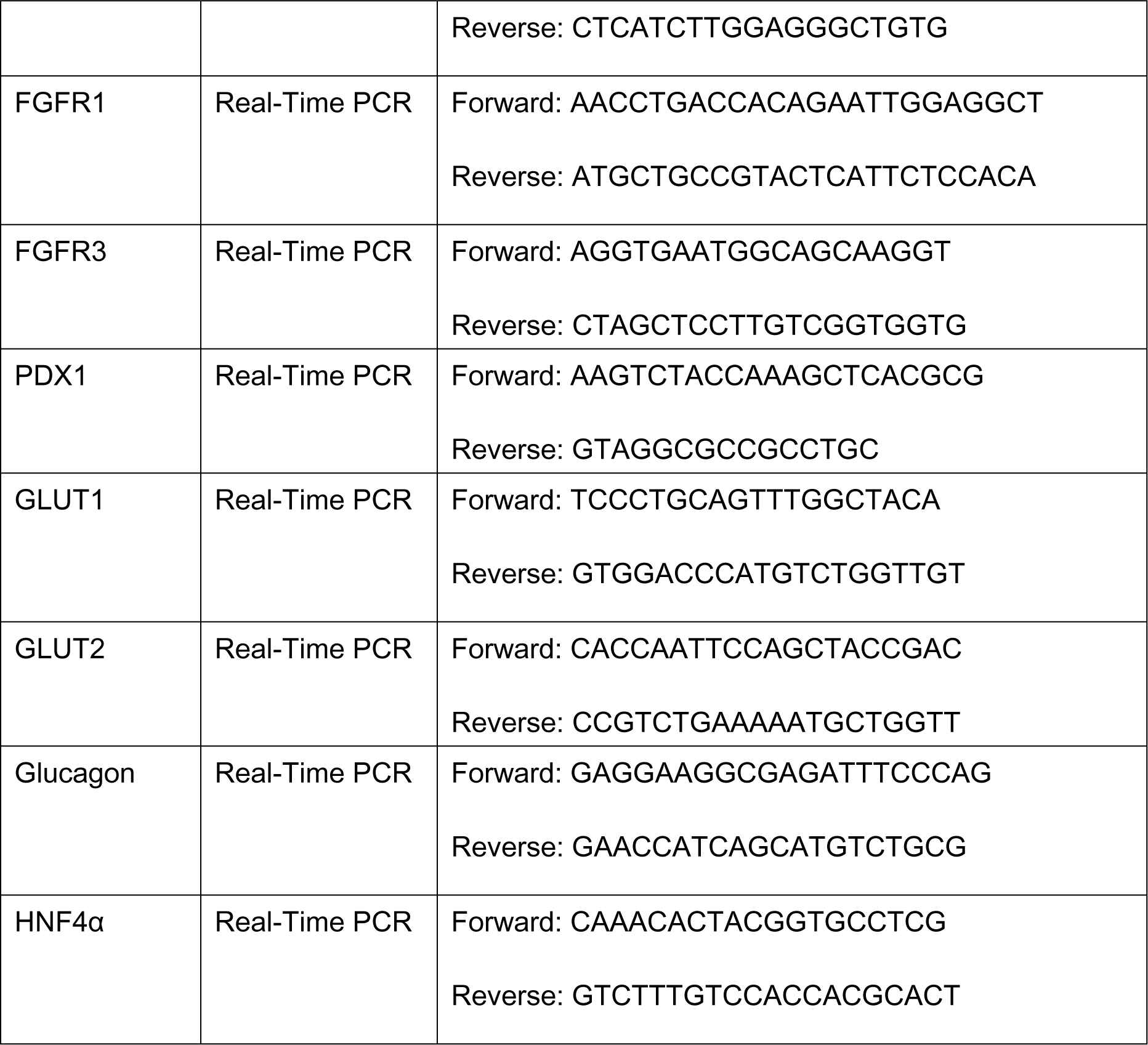
PCR and qPCR primers

**Table S4:** Antibodies for flow cytometry and immunofluorescence analysis

|  |  |
| --- | --- |
| PE Mouse anti-Human Sox17 | BD Biosciences |
| Alexa Fluor® 488 Mouse anti-PDX-1 | BD Biosciences |
| PE Mouse Anti-Nkx6.1 | BD Biosciences |
| Alexa Fluor® 647 Mouse Anti-Nkx6.1 | BD Biosciences |
| Insulin (C27C9) Rabbit mAb (Alexa Fluor® 488 Conjugate) | Cell Signaling Technology |
| APC anti-Human CD26 | Biolegend |
| PE Mouse Anti-Human CD49a | BD Biosciences |
| Anti-Tra-1-60-PE, human | MACS Miltenyi Biotec |
| Alexa Fluor® 647 Mouse IgG1 κ Isotype Control | BD Biosciences |
| FITC Mouse IgG2a, κ Isotype Control | BD Biosciences |
| PE Mouse IgG1, κ Isotype Control | BD Biosciences |
| APC Mouse IgG2a, κ Isotype Control | Biolegend |
| Rabbit IgG Isotype Control (Alexa Fluor® 488 Conjugate) | Cell Signaling Technology |
| FITC Annexin V | BD Biosciences |
| PDX1 Mouse Monoclonal Antibody | Origene |
| Homeobox protein Nkx-6.1 | Developmental Studies |
| Purified Mouse Anti-Ki-67 | BD Biosciences |
| Anti-KCNQ1 antibody | ATLAS ANTIBODIES |
| FLEX Polyclonal Guinea Pig Anti-Insulin Ready-to-use | Agilent |
| Insulin (C27C9) Rabbit mAb | Cell signaling Technology |
| Monoclonal Anti-Glucagon antibody | Sigma |
| C-Peptide Antibody | Cell signaling Technology |
| Sox2 (L1D6A2) Mouse mAb | Cell signaling Technology |
| Oct-4 Antibody | Cell signaling Technology |
| Actin, pan Ab-5 | Dianova |
| Alexa Fluor™ 488 Goat Anti-Mouse | Invitroge |
| Goat anti-Rabbit IgG (H+L) Highly Cross-Adsorbed<br>Secondary Antibody, Alexa Fluor 555 | ThermoFisher |
| Goat anti-Guinea Pig IgG (H+L) Highly<br>Cross-Adsorbed Secondary antibody, Alexa Fluor 647 | ThermoFisher |
| HPR-Anti Rabbit IgG (H+L) | Thermo Scientific |
| HPR-Anti Mouse IgG (H+L) | Thermo Scientific |

